# Uncovering the hidden antibiotic potential of Cannabis

**DOI:** 10.1101/833392

**Authors:** Maya A. Farha, Omar M. El-Halfawy, Robert T. Gale, Craig R. MacNair, Lindsey A. Carfrae, Xiong Zhang, Nicholas G. Jentsch, Jakob Magolan, Eric D. Brown

**Author notes:** These authors contributed equally (Maya A. Farha, Omar M. El-Halfawy).

## Abstract

The spread of antimicrobial resistance continues to be a priority health concern worldwide, necessitating exploration of alternative therapies. *Cannabis sativa* has long been known to contain antibacterial cannabinoids, but their potential to address antibiotic resistance has only been superficially investigated. Here, we show that cannabinoids exhibit antibacterial activity against MRSA, inhibit its ability to form biofilms and eradicate pre-formed biofilms and stationary phase cells persistent to antibiotics. We show that the mechanism of action of cannabigerol is through targeting the cytoplasmic membrane of Gram-positive bacteria and demonstrate *in vivo* efficacy of cannabigerol in a murine systemic infection model caused by MRSA. We also show that cannabinoids are effective against Gram-negative organisms whose outer membrane is permeabilized, where cannabigerol acts on the inner membrane. Finally, we demonstrate that cannabinoids work in combination with polymyxin B against multi-drug resistant Gram-negative pathogens, revealing the broad-spectrum therapeutic potential for cannabinoids.

---

Public Health agencies around the globe have identified antimicrobial resistance as one of the most critical challenges of our time. The rapid and global spread of antimicrobial-resistant organisms in recent years has been unprecedented. So much so that the world health organization (WHO) published its first ever list of antibiotic-resistant “priority pathogens”, made up of 12 families of bacteria that pose the greatest threat to human health^1^. Among them, *Staphylococcus aureus* is the leading cause of both healthcare and community-associated infections worldwide and a major cause for morbidity and mortality^2^, especially with the emergence and rapid spread of methicillin-resistant *S. aureus* (MRSA), which is resistant to all known β-lactam antibiotics^3^. Worse yet, resistance to vancomycin, linezolid and daptomycin has already been reported in MRSA clinical strains, compromising the therapeutic alternatives for life-threatening MRSA infections^4^. Further, antibiotic-resistant Gram-negative infections have increasingly become a pressing issue in the clinic. Indeed, of the bacteria highlighted by the WHO, 75% are Gram-negative organisms. Among the currently approved antibiotics in clinical use, the latest discovery of a new drug class dates back to more than 30 years ago. The rapid loss of antibiotic effectiveness and diminishing pipeline beg for the exploration of alternative therapies.

*Cannabis* plants are important herbaceous species that have been used in folk medicine for centuries. Increasing scientific evidence is accumulating for the efficacy of its metabolites in the treatment, for example, of epilepsy, Parkinson disease, analgesia, multiple sclerosis, Tourette’s syndrome and other neurological diseases^5^. At a very nascent stage are investigations into the potential of cannabis metabolites as antibacterial therapies. To date, assessments of their antibacterial activity have been few and superficial. *In vitro* studies have shown that cannabinoids inhibit the growth of Gram-positive bacteria, mostly *S. aureus*, with no detectable activity against Gram-negative organisms^6-9^, where the clinical need is highest. Further, the mechanism of action has remained elusive and there has been little validation of antibacterial activity *in vivo*.

Here, we show that cannabinoids exhibit antibacterial activity against MRSA, inhibit its ability to form biofilms, eradicate pre-formed biofilms and stationary phase cells persistent to antibiotics. We reveal that the mechanism of action of cannabigerol (CBG) is through targeting the cytoplasmic membrane of Gram-positive bacteria and demonstrate *in vivo* efficacy of CBG in a murine systemic infection model caused by MRSA. We also show that cannabinoids are effective against Gram-negative organisms whose outer membrane is permeabilized, where CBG acts on the inner membrane. Finally, we demonstrate that cannabinoids work in combination with polymyxin B against multi-drug resistant Gram-negative pathogens, revealing the broad-spectrum therapeutic potential for cannabinoids. In all, our findings position cannabinoids as promising leads for antibacterial development that warrant further study and optimization.

## Results and Discussion

We began our study investigating the antibacterial, anti-biofilm and anti-persister activity of a variety of commercially available cannabinoids, including the five major cannabinoids, cannabichromene (CBC), cannabidiol (CBD), cannabigerol (CBG), cannabinol (CBN), and Δ^9^-tetrahydrocannabinol (THC), as well as a selection of their carboxylic precursors (pre-cannabinoids) and other synthetic isomers (18 unique molecules total, Fig. 1) against methicillin-resistant *S. aureus* (MRSA) (Supplementary Table 1). Susceptibility tests were conducted according to the Clinical and Laboratory Standards Institute (CLSI) protocol against MRSA USA300, a highly virulent and prevalent community-associated MRSA. Overall, antibacterial activities for the five major cannabinoids (and some of their synthetic derivatives) were in line with previously published work^6-8^. Seven molecules were potent antibiotics with minimum inhibitory concentration (MIC) values of 2 µg/mL, including CBG, CBD, CBN, cannabichromenic acid (CBCA) and THC along with its Δ^8^- and exo-olefin regioisomers. We observed moderate loss of potency associated with the benzoic acid moiety (CBG, CBD, and THC were more potent than CBGA, CBDA, THCA) and when n-pentyl substituent was replaced with n-propyl (CBD and THC were superior to CBDV and THCV) (Supplementary Table 1). These two modifications appeared to have an additive detrimental effect on antibacterial activity (THCVA, CBDVA). The two most common human metabolites of THC, (±) 11-nor-9-carboxy-Δ^9^-THC, and (±) 11-hydroxy-Δ^9^-THC, as well as cannabicylol were inactive at the highest concentrations screened (MIC > 32 µg/mL) (Supplementary Table 1).

**Fig. 1.**
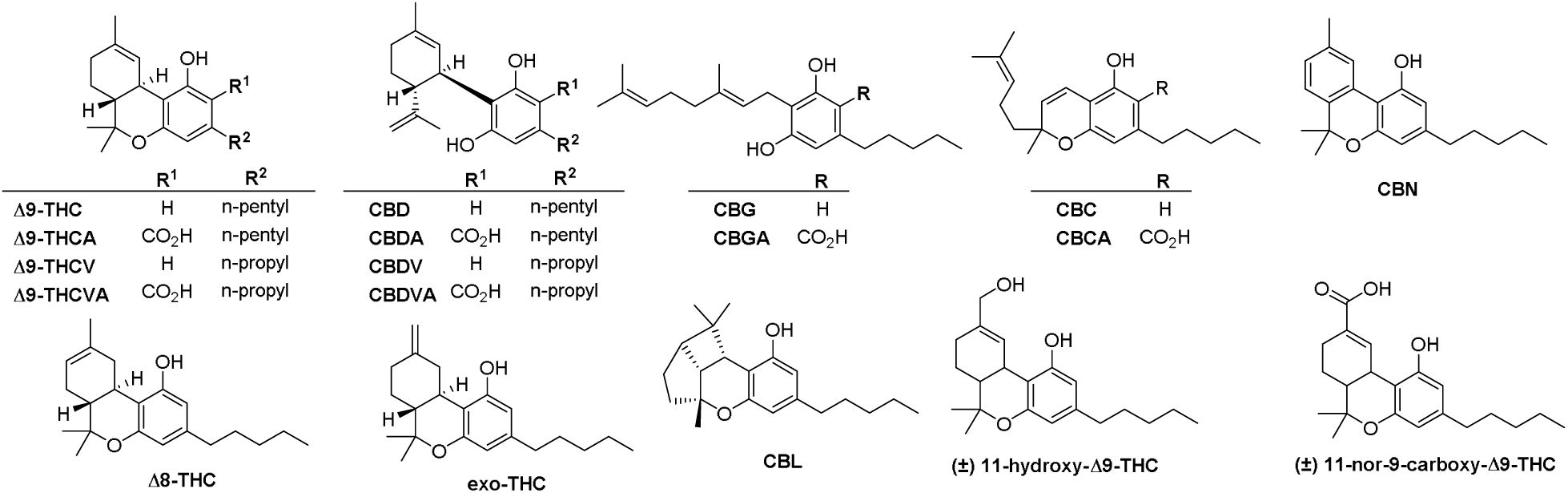
Structures of the cannabinoids surveyed in this study.

Biofilm formation by MRSA, typically on necrotic tissues and medical devices, is considered an important virulence factor influencing its persistence in both the environment and the host organism^10^. These highly structured surface-associated communities of MRSA are typically associated with increased resistance to antimicrobial compounds and are generally less susceptible to host immune factors. We assessed the ability of the various cannabinoids to inhibit the formation of biofilms by MRSA, using static abiotic solid-surface assays in which MRSA was treated with increasing concentrations of cannabinoids under conditions favouring biofilm formation (Supplementary Fig. 1). In all, the degree of inhibition of biofilm formation correlated with antibacterial activity; those cannabinoids with potent activity against MRSA strongly suppressed biofilm formation and vice versa (Supplementary Fig. 1, Supplementary Table 1). The five major cannabinoids clearly repressed MRSA biofilm formation, with CBG (Fig. 2a) exhibiting the most potent anti-biofilm activity. Indeed, as little as 0.5 μg/mL (1/4 MIC) of CBG inhibited biofilm formation by ⍰50% (Fig. 2b). Thus, this experiment underlined the strong inhibitory effect of cannabinoids on biofilm formation; this sub-MIC level of CBG did not affect planktonic growth (Supplementary Fig. 2). Interestingly, we also evaluated the effect of CBG on pre-formed biofilms by determining its minimal biofilm eradication concentration (MBEC); CBG could eradicate pre-formed biofilms of MRSA USA300 at 4 µg/mL (Fig. 2c).

**Fig. 2.**
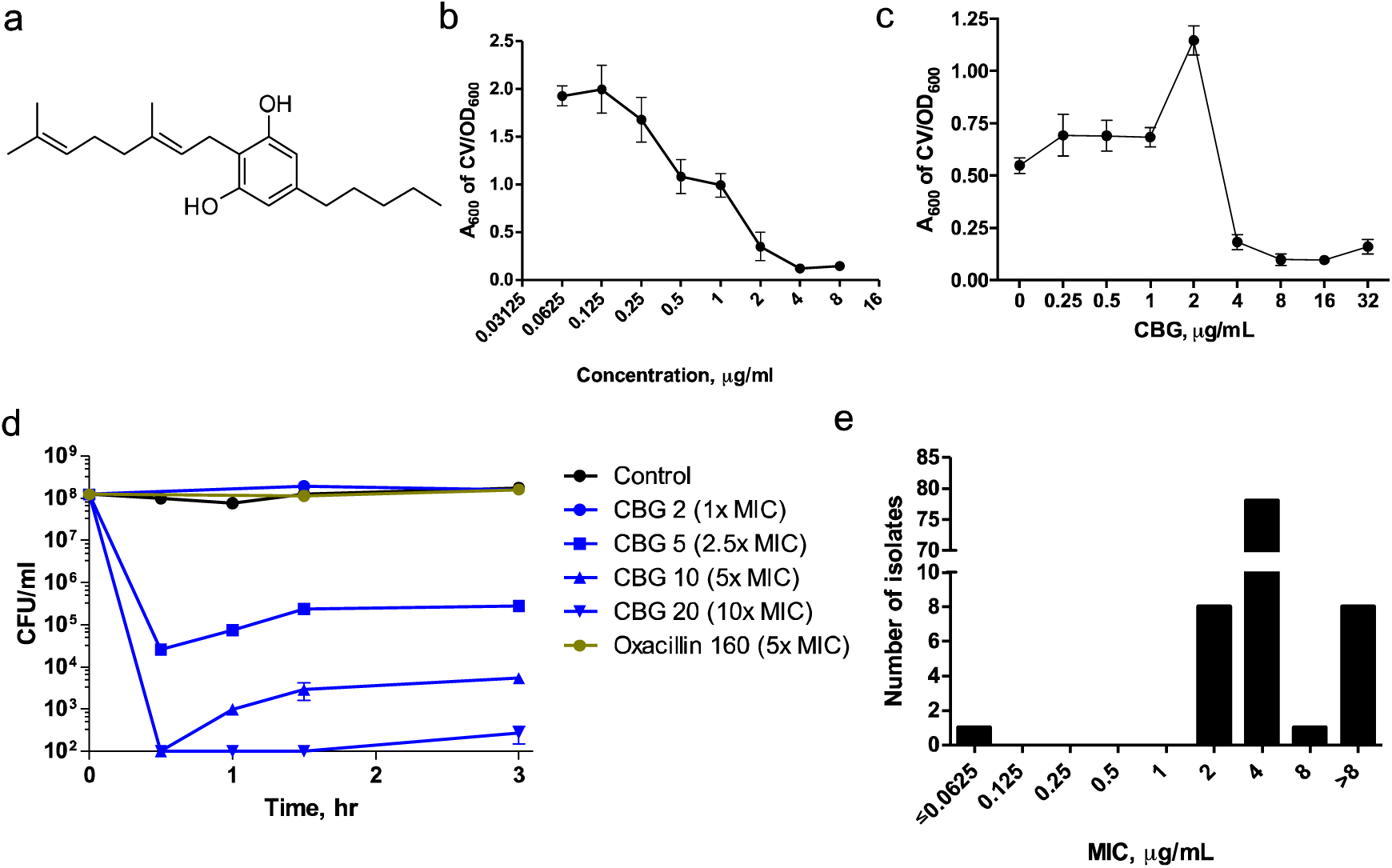
Cannabigerol (CBG) is a potent antibacterial, anti-biofilm and anti-persister cannabinoid. **a**, Chemical structure of CBG **b**, Concentration dependence for inhibition of MRSA biofilm formation by CBG. Shown is the average A_600nm_ measurements of crystal violet stained biofilms and normalized by the OD_600_ of planktonic cells with error bars representing one standard error of the mean, S.E.M. (n□= □4). c, Minimum biofilm eradication concentration of CBG. CBG is capable of eradicating pre-formed biofilms at a concentration of 4µg/mL (n=8).**d**, Time-kill curve of *S. aureus* USA300 persisters by CBG compared to oxacillin shown as mean ±S.E.M (n=4). CBG rapidly eradicated a population of ∼10^8^ CFU/ml MRSA persisters to below the detection threshold within 30 minutes of treatment. On the other hand, the β-lactam oxacillin at 160 µg/mL (5x MIC) did not show any activity against the same population of persisters. **e**, MIC_90_ distribution of CBG against clinical isolates of MRSA (n=96). The MIC_90_ is 4 µg/mL.

Another challenge in the treatment of MRSA infections is the formation of non-growing, dormant ‘persister’ subpopulations that exhibit high levels of tolerance to antibiotics^11-13^. Persister cells have a role in chronic and relapsing *S. aureus* infections^14^ such as osteomyelitis^15^, and endocarditis^16^. Here, we evaluated the killing activity of a series of cannabinoids against persisters derived from stationary phase cells of MRSA USA300 (Supplementary Fig. 3). These have been previously shown to be tolerant to conventional antibiotics such as gentamicin, ciprofloxacin and vancomycin^11, 17-18^. In general, the anti-persister activity correlated with potency against actively dividing cells as determined by MIC assays (Supplementary Table 1). Again, CBG was the most potent cannabinoid against persisters, whereas oxacillin and vancomycin were ineffective at concentrations that otherwise kill actively dividing cells (Supplementary Fig. 3, Fig. 2d). More specifically, CBG killed persisters in a concentration-dependent manner starting at 5 µg/ml. Notably, CBG rapidly eradicated a population of ∼10^8^ CFU/ml MRSA persisters to below the detection threshold within 30 minutes of treatment (Fig. 2d).

We selected CBG (Fig.2a) for further studies of mechanism and *in vivo* efficacy. Not only did CBG potently inhibit MRSA (MIC 2 µg/mL), repress biofilm formation (Fig. 2b), eradicate pre-formed biofilms (Fig. 2c) and effectively eradicate persister cells (Fig. 2d), but it is non-psychotropic, non-sedative and constitutes a component of *Cannabis* for which there is high therapeutic interest^19^. While it has many desirable pharmacological properties, CBG also possesses several desirable physico-chemical properties as a medicinal chemistry lead in terms of molecular weight, number of hydrogen donors and acceptors, number of rings and rotatable bonds (Table 1). Nevertheless, CBG suffers from high lipophilicity (high log P) and low aqueous solubility (Table 1). These values were not a bottleneck to our studies, but in moving CBG as a lead, such properties could be addressed in medicinal chemistry campaigns. To our advantage, we were able to synthesize CBG efficiently from olivetol and geraniol, two inexpensive precursors, in one synthetic operation. We were cognisant that such facile synthetic access would enhance the potential for subsequent medicinal chemistry-based development efforts. We determined the MIC_90_ of CBG against 96 clinical isolates of MRSA using the CLSI protocol. The corresponding frequency distribution of MICs is presented in Fig. 2e. The MICs ranged from 2 – 8 µg/mL with a resulting MIC_90_ of 4 µg/mL. The range of MIC values was tight with only one outlier strain detected to have a very low MIC of 0.0625. Overall, this activity compares favourably with conventional antibiotics for these multi-drug resistant strains.

**Table 1.** Physiochemical properties of CBG as calculated by ACD/Percepta software.

| Phys Chem Profile |  | Lead-like? |
| --- | --- | --- |
| MW | 316.48 | good |
| Log P | 6.74 | very lipophilic |
| H-Donors | 2 | good |
| H-Acceptors | 2 | good |
| Rotatable Bonds | 9 | good |
| Rings | 1 | good |
| Predicted solubility | 0.0003 mg/mL | highly insoluble |

Given its growth inhibitory action on Gram-positive bacteria, we reasoned isolating resistant mutants to CBG would be a straightforward approach to gather insights into its bacterial target. Indeed, resistance mutations can often be mapped to a drug’s molecular target^20^. To this end, MRSA was repeatedly challenged with various lethal concentrations of CBG, ranging from 2-16x MIC, to select for spontaneous resistance in MRSA (Fig. 3). No spontaneously resistant mutants were obtained, indicating a frequency of resistance less than 10^−10^ for MRSA. We also attempted to allow MRSA bacteria to develop resistance to CBG by sequential subcultures via 15-day serial passage in liquid culture containing sub-MIC concentrations of CBG and, again, no change in the MIC of CBG was detected (Fig. 3). While these experiments were unsuccessful probes of mechanism, they suggested very low rates of resistance for CBG, a highly desirable property for an antibiotic.

**Fig. 3.**
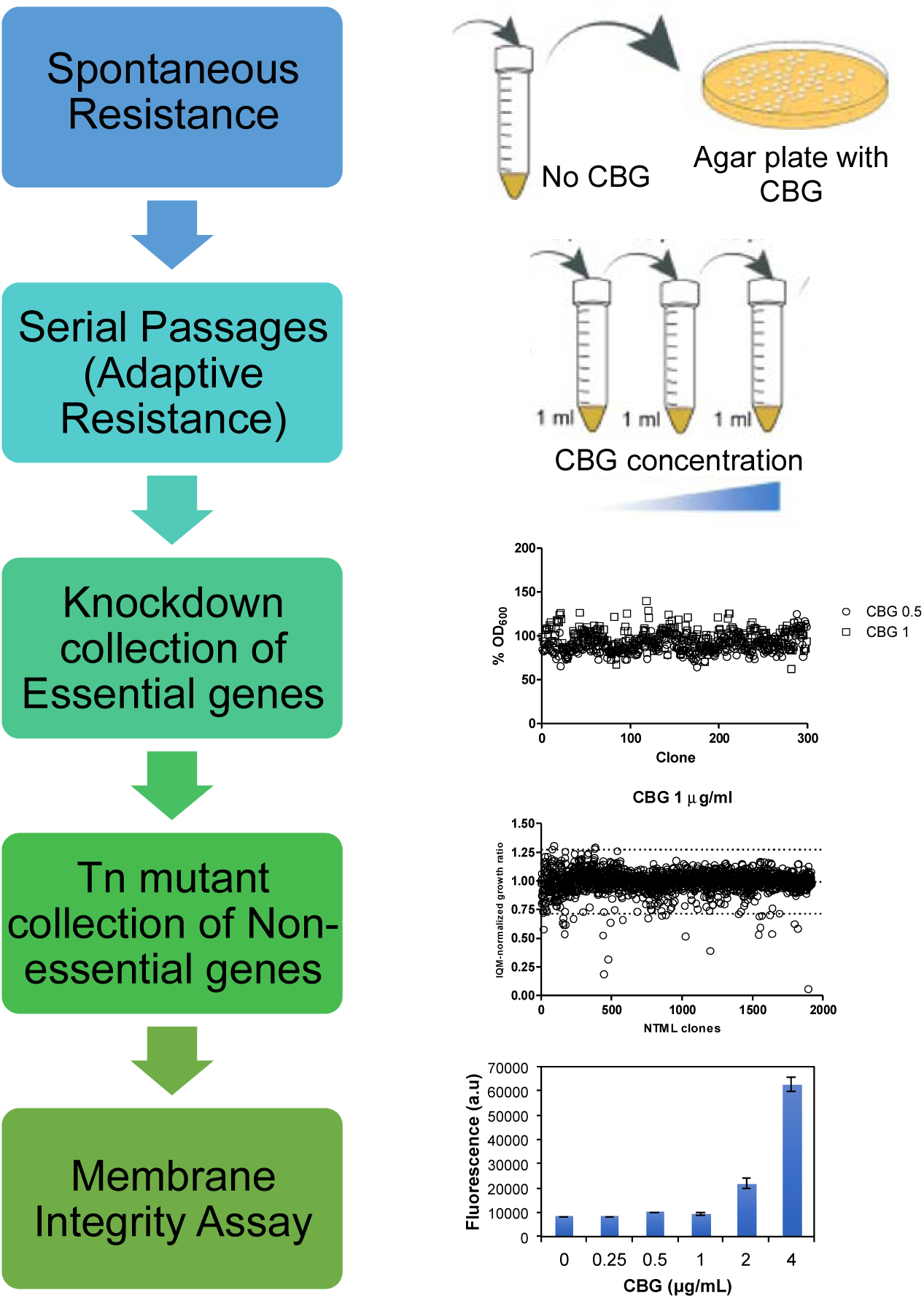
CBG is active on the cytoplasmic membrane of MRSA. Overview of strategies for mechanism of action determination, culminating in the finding that CBG is active on the cytoplasmic membrane, as determined by dose-dependent increases in DiSC_3_(5) fluorescence.

We turned to chemical genomic analysis to generate hypotheses for the target of CBG. Such studies can reveal patterns of sensitivity among genetic loci that are characteristic of the mechanism of action of an antibacterial compound^21^. We confirmed that the model Gram-positive bacterium *B. subtilis* was susceptible to CBG (MIC 2 µg/mL), and screened a CRISPR interference knockdown library of all essential genes in *B. subtilis*^*22*^ for further sensitization to CBG. In the absence of induction, relying on basal repression (which leads to a ∼3-fold repression of the knockdown library^*22*^), we were unable to detect any knockdowns sensitized to sub-lethal concentrations of CBG (Fig. 3). Low-level induction identified some sensitive and some suppressing clones, however follow-on work with the individual knockdowns in liquid culture via full checkerboard analysis (combining xylose, the inducer, with CBG) failed to confirm sensitivity or suppression. In all, we were unable to identify *bona fide* chemical genetic interactions among essential genes of *B. subtilis* and CBG. We next aimed to query the non-essential gene subset, this time using the Nebraska Transposon Mutant Library, a sequence-defined transposon mutant library consisting of 1,920 strains, each containing a single mutation within a nonessential gene of CA-MRSA USA300^23^, again looking for genetic enhancers or suppressors to generate target hypotheses (Fig. 3, Supplementary Fig. 4a). While we were unable to uncover genetic suppressors at supra-lethal concentrations of CBG, we identified 41 transposons as sensitive across 3 different sub-lethal concentrations of CBG (Supplementary Table 2). Analysis of these transposons revealed a significant enrichment for genes encoding proteins that are localized at the cytoplasmic membrane (Supplementary Fig. 4b) and enrichment for genes encoding functions in processes that take place at the cytoplasmic membrane, such as cellular respiration and electron transport chain (Supplementary Fig. 4c). In all, chemical genomic profiling with CBG generally linked its activity to cytoplasmic membrane function.

The lack of clear targets among the essential gene products, the predominance of chemical genetic interactions linked to membrane function, and the difficulty generating resistant mutants, suggested that CBG might act on the cytoplasmic membrane of MRSA. Indeed, the propensity of membrane-active compounds to generate resistance is frequently low^24^. Further, the bacterial membrane is critical for cell function and survival, and is essential irrespective of the metabolic status of the cell, including non-growing and persisting cells^24^. The strong action of CBG on persister cells would be consistent with such a mode of action. Thus, we assessed the ability of CBG to disrupt membrane function using the membrane potential-sensitive probe, 3,3’-dipropylthiadicarbocyanine iodide (DiSC_3_(5)). In DiSC_3_-loaded MRSA cells, CBG caused a dose-dependent increase in fluorescence that occurred at a concentration consistent with the MIC of CBG (Fig. 3). To probe the possibility that CBG selectively dissipated membrane potential (ΔΨ) component of proton motive force, we tested for synergy with sodium bicarbonate, a known perturbant of ΔpH, that has been shown to synergize with molecules that reduce ΔΨ^25^. A lack of synergy between these compounds suggested CBG disrupts the integrity of the cytoplasmic membrane (Supplementary Fig. 5). In order to evaluate the potential membrane activity of CBG on mammalian cells, we evaluated its hemolytic toxicity across varying concentrations (Supplementary Fig. 6). CBG was hemolytic only at 32 µg/mL, above its MIC of 2 µg/mL, suggesting some specificity for prokaryotic cells.

Having established strong *in vitro* potency for CBG against MRSA, we next sought to evaluate the *in vivo* efficacy in a murine systemic infection model of MRSA. The effect of CBG on a systemic infection mediated by the CA-MRSA USA300 strain is shown in Fig. 4. A dose-dependence study (Supplementary Fig. 7) informed that a dose of 100 mg/kg is most effective while remaining tolerable. To evaluate tolerability, we treated mice with 100 mg/kg CBG and assessed their change in body weight over various time points and found no significant changes (Supplementary Fig. 8). Additionally, no signs of acute toxicity have been reported in a pharmacokinetic study of 120-mg/kg doses of CBG^26^.. Overall, CBG displayed a significant reduction in bacterial burden in the spleen by a factor of 2.8-log_10_ in CFU compared to the bacterial titer seen with the vehicle (*p*□<□0.001, Mann–Whitney *U*-test). Further, the *in vivo* efficacy of CBG was comparable to that of the antibiotic control, vancomycin administered at a similar dose. In all, CBG displayed promising levels of efficacy in the systemic infection model.

**Fig. 4.**
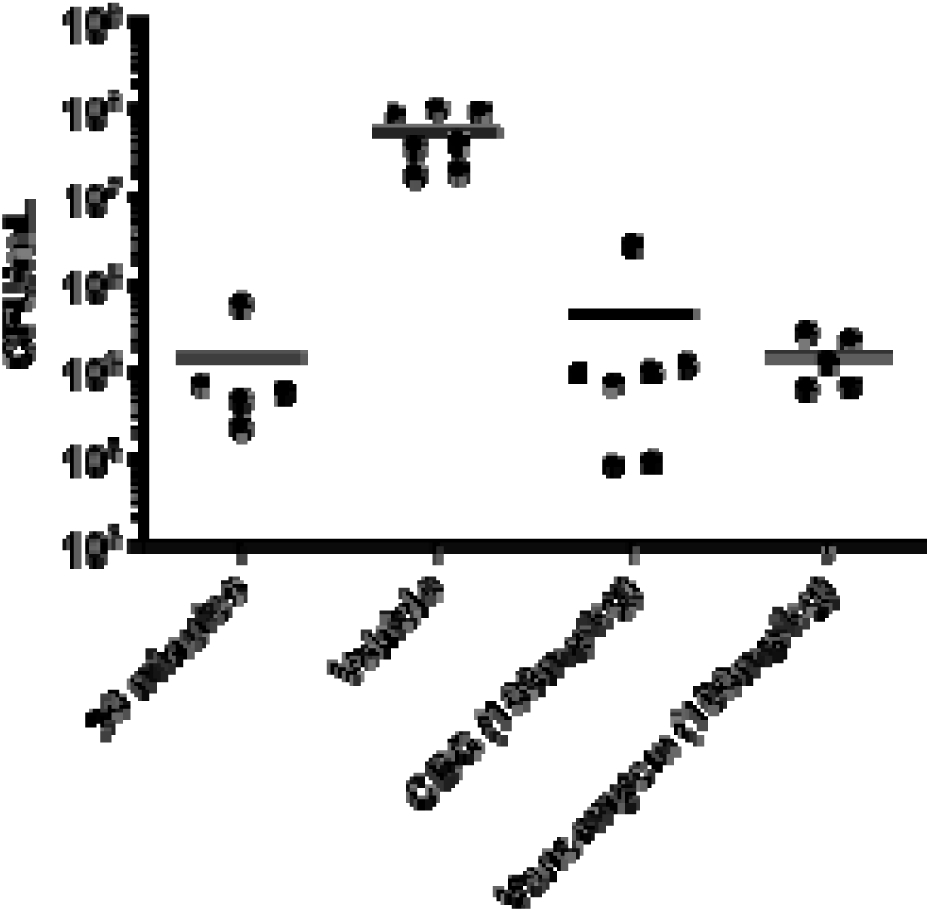
CBG is efficacious in a systemic mouse model of *S. aureus* infection. A single-dose treatment is administered immediately post-infection: vehicle control (n=7, i.p.), CBG (n=7, 100 mg□kg^−1^, i.p.) or vancomycin control (n=5, 100 mg□kg^−1^, i.p.). Colony-forming units (CFU) within spleen tissue were enumerated at 7 h post-infection. CFUs within spleen tissue were also enumerated at an early time point of 30 minutes post-infection following vehicle control treatment (n=5, i.p.). Horizontal lines represent the geometric mean of the bacterial load for each treatment group. Administration of CBG resulted in a 2.8-log_10_ reduction (*p*□< □0.001, Mann–Whitney *U*-test) in CFU when compared to the vehicle control.

To date, antibacterial activity of cannabinoids against Gram-negative organisms has largely been ruled out, since reported MICs values fall in the 100-200 µg/mL range^7-8^. We confirmed this, obtaining MICs >128 µg/mL for all of the tested cannabinoids against the model Gram-negative organism *Escherichia coli*. Given the observed action of CBG on the cytoplasmic membrane of MRSA, we reasoned that CBG (and other cannabinoids) might be equally effective on the Gram-negative counterpart, the inner membrane. Further, just as many antibacterial compounds fail to work against Gram-negative pathogens due to a permeability barrier^27^, we reasoned that low permeability across the outer membrane (OM) may be the reason for the poor efficacy of cannabinoids. Thus, we investigated the antibacterial profile of the five major cannabinoids against *E. coli*, where their permeation was facilitated through the OM by means of chemical perturbation. To this end, we set up checkerboard assays to assess the interaction of CBG (Fig. 5a) and the four other main cannabinoids (Supplementary Fig. 9) with the membrane perturbant, polymyxin B against *E. coli*. Remarkably, all five major cannabinoids gained potent activity in the presence of sub-lethal concentrations of polymyxin B. Indeed, all interactions were deemed synergistic (Fig. 5, Supplementary Fig. 9). For example, CBG, which was inactive against *E. coli* (>128 µg/mL), was strongly potentiated when combined with a sub-lethal concentration of polymyxin B (1 µg/mL in the presence of 0.062 µg/mL polymyxin B). A similar synergistic interaction was observed with polymyxin B nonapeptide, a less toxic derivative of polymyxin B that strictly perturbs the outer membrane in Gram-negative bacteria (Supplementary Fig. 10), suggesting that induction of outer membrane permeability is sufficient to allow entry and activity of CBG. We further assessed whether OM perturbation by genetic means would lead to similar results by evaluating the activity of CBG against a number of strains where the OM was compromised (Fig. 5b). In an *E. coli* Δ*bamB*Δ*tolC* deletion strain, which renders *E. coli* hyperpermeable to many small molecules, due to loss of BamB, a component of the β-barrel assembly machinery for OM proteins and TolC, the efflux channel in the outer membrane, CBG had a MIC of 4 µg/mL, on par with its Gram-positive activity. Similarly, in a hyperporinated, Δ9 strain of *E. coli*, where a recombinant pore was introduced in the OM and all nine known TolC-dependent transporters deleted^28^, CBG activity became evident with a MIC of 8 µg/mL. Finally, in an *Acinetobacter baumannii* deficient in lipooligosaccharide (LOS-), which effectively alters the permeability of the OM^29^, CBG activity was enhanced greater than 128-fold, resulting in a MIC value of 0.5 µg/mL. Overall, these results suggest that cannabinoids face a permeability barrier in Gram-negative bacteria and further imply that cannabinoids inhibit a bacterial process present in Gram-negative pathogens, and likely common to that in Gram-positive pathogens.

**Fig. 5.**
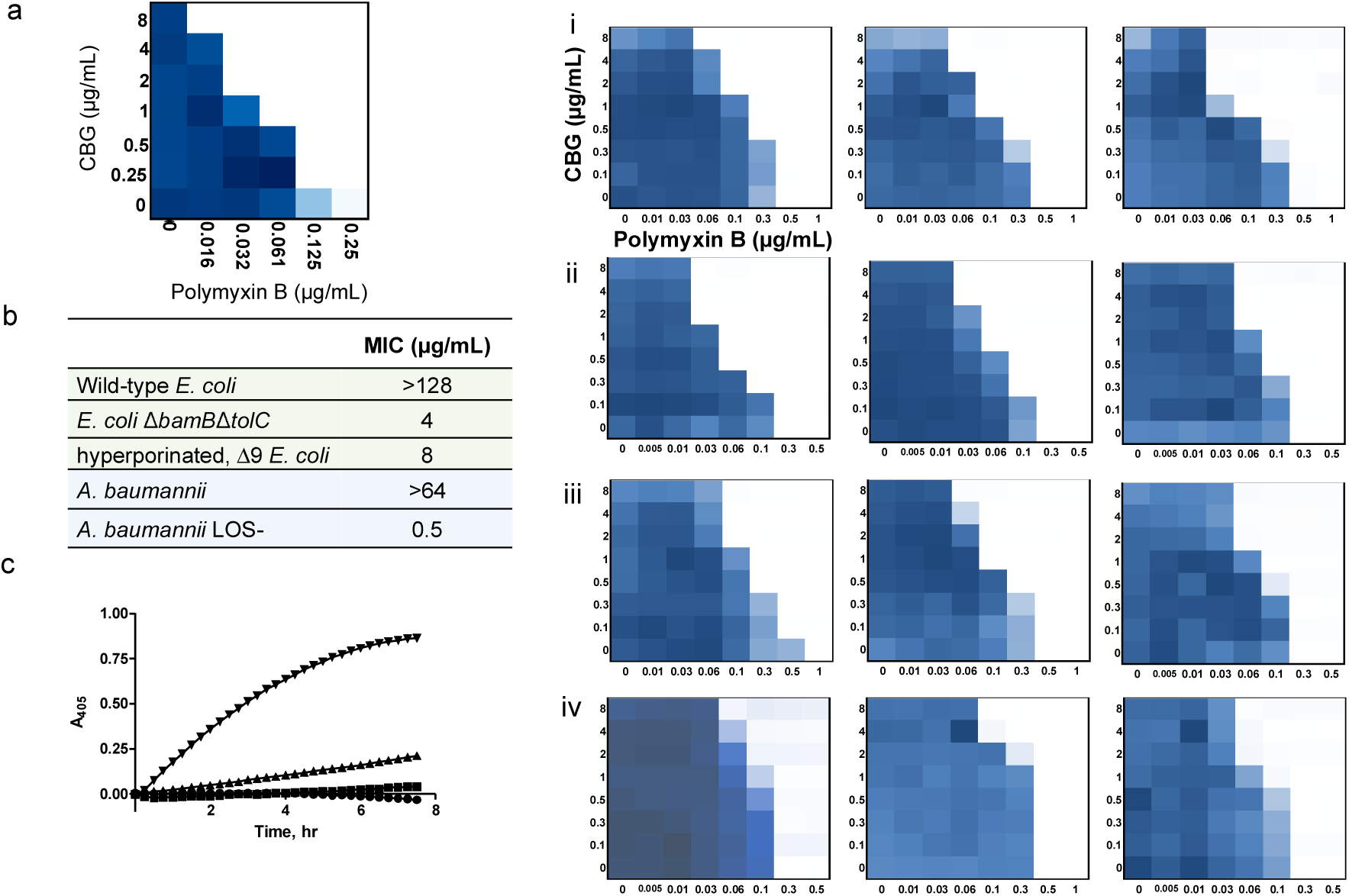
CBG is active against Gram-negative bacteria whose outer membrane is permeabilized, where it acts on the inner membrane. **a**, Checkerboard analysis of CBG in combination with polymyxin B against *E. coli*. The extent of inhibition is shown as a heat plot, such that the darkest blue color represents full bacterial growth. **b**, CBG becomes active against Gram-negative bacteria in various genetic backgrounds where the outer membrane is compromised. **c**, CBG acts on the IM of *E. coli* but only in the presence of sub-lethal concentration of polymyxin B (PmB), unmasking cytoplasmic β-galactosidase leading to hydrolysis of ONPG as detected via absorbance reads at 405 nm over time. Conditions were as follows: control (circles), CBG 2 µg/mL (squares), PmB 0.125 µg/mL (triangles) and PmB 0.125 µg/mL + CBG 2 µg/mL (inverted triangles). **d**, CBG in combination with polymyxin B against multi-drug resistant clinical isolates of i, *A. baumannii*, ii, *E. coli*, iii, *K. pneumoniae*, iv, *P. aeruginosa*. The extent of inhibition is shown as a heat plot, such that the darkest blue color represents full bacterial growth.

To this end, we investigated whether CBG acted on the inner membrane (IM) of *E. coli* as well as the OM. IM and OM permeability were determined, respectively, from ortho-Nitrophenyl-β-galactoside (ONPG) and nitrocefin hydrolysis in an *E. coli* strain constitutively expressing a cytoplasmic β-galactosidase and a periplasmic β-lactamase while lacking the lactose permease, as described in the literature^30^. As shown in Fig. 5c, CBG specifically acted on the IM, and only in the presence of polymyxin B at a sub-lethal concentration that had minimal effects on the IM alone. We observed that CBG (+polymyxin B) induced major permeability changes in the inner membrane, indicated by a time-dependent marked increase in optical density values due to ONPG hydrolysis as a result of unmasking the cytoplasmic β-galactosidase, which can only occur with destabilization of IM (Fig. 5c). CBG exhibited no action on the OM (Supplementary Fig. 11). Overall, the mechanism of bacterial killing by CBG in *E. coli* is likely loss of IM integrity and requires antecedent OM permeabilization.

Combination antibiotic therapy is becoming an increasingly attractive approach to combat resistance^31^. So too is the strategy of using an OM perturbing molecule to facilitate the permeation of compounds that are otherwise active only on Gram-positive bacteria^32^. We assessed the therapeutic potential of the adjuvant polymyxin B in combination with CBG to inhibit the growth of priority Gram-negative pathogens such as *A. baumannii, E. coli, Klebsiella pneumoniae*, and *Pseudomonas aeruginosa* (Fig. 5d). We employed conventional checkerboard assays to determine the interaction and potency of CBG and polymyxin B when used concurrently against various multi-drug resistant clinical isolates. In all cases, synergy was evident, suggesting the potential for combination therapy of the cannabinoids with polymyxin B against Gram-negative bacteria.

In summary, we have investigated the therapeutic potential of cannabinoids, specifically CBG, through a comprehensive study of *in vitro* potency on biofilms and persisters, as well as mechanism of action studies and *in vivo* efficacy experiments. Most notably, we have uncovered the hidden broad-spectrum antibacterial activity of cannabinoids and demonstrated the potential of CBG against Gram-negative priority pathogens. Taken together, our findings lend credence to the idea that cannabinoids may be produced by *Cannabis sativa* as a natural defense against plant pathogens. Notwithstanding, cannabinoids are well-established as drug compounds that have favourable pharmacological properties in humans. The work presented here suggests that the cannabinoid chemotype represents an attractive lead for new antibiotic drugs.

## Acknowledgements

This work was supported by a salary award to E.D.B from the Canada Research Chairs program and operating funds to E.D.B from a CIHR Foundation grant (FDN-143215); by a Michael G. DeGroote Centre for Medicinal Cannabis Research post-doctoral fellowship to O.M.E. Synthetic chemistry was supported by McMaster’s Faculty of Health Sciences and the Michael G. DeGroote Institute for Infectious Disease Research. We thank Shawn French for preparing the graphical abstract.

## Author contributions

M.A.F., O.M.E., R.T.G., and E.D.B. conceived and designed the research. M.A.F. and O.M.E., performed all experiments and analyzed data with the exception of the mouse infection model and the synthesis of CBG. C.R.M. and L.A.C. performed the mouse infection model. X.Z. and N.G.J. optimized a scalable synthesis of CBG, supervised by J.M. M.A.F. and E.D.B. wrote the paper, with large input from O.M.E. All authors approved the final version.

## METHODS

### Strains and reagents

Supplemental Table 3 lists bacteria and plasmids used in this work. Supporting Information *S. aureus* Table lists the 96 clinical isolates, along with body site location and year of isolation. Bacteria were grown in cation-adjusted Mueller Hinton broth (CAMHB) at 37°C, unless otherwise stated. Antibiotics were obtained from Sigma, Oakville, ON, Canada. Pure cannabinoid solutions purchased from Sigma, Oakville, ON, Canada were used for all experiments. Only CBG was synthesized in larger quantity as described below.

### Antimicrobial susceptibility testing

Minimum inhibitory concentration (MIC) determination and checkerboard assays were conducted following the guidelines of CLSI for MIC testing by broth microdilution^33^. Persister killing activity of cannabinoids was evaluated against stationary-phase cells of *S. aureus* as previously described^35^.

### *B. subtilis* CRISPRi essential gene knockdown strain collection screen

Overnight cultures of the collection^22^ (at a 96-well density, *n* = 289) were performed using the Singer rotor HDA (Singer Instruments, United Kingdom) in CAMHB. Subsequently, CAMHB with or without CBG were inoculated using the singer rotor at 96-well density. These experiments were performed either in the presence of 0.05% xylose (allowing low level of *dcas9* expression) or with no xylose induction (basal *dcas9* expression). The plates were incubated at 37°C and OD_600_ was read after 24 h.

### *S. aureus* Nebraska Transposon Mutant Library (NTML) screen

Overnight cultures of the NTML^23^ (at a 384-well density) were performed using the Singer rotor HDA (Singer Instruments, United Kingdom) in CAMHB containing erythromycin (5 µg/mL). Subsequently, CAMHB with or without CBG were inoculated using the singer rotor at 384-well density. The plates were incubated at 37°C and OD_600_ was read after 24 h. Cellular localization and functional (gene ontology, GO-term) enrichment analyses were performed using Pathway Tools software and MetaCyc database^36^.

### Selection of suppressor mutants of CBG activity in *S. aureus*

Spontaneous suppressor mutants were selected for in liquid culture. Briefly, isolated colonies were resuspended in PBS and diluted to a final OD_600_ of 0.05 into 200 μL of CAMHB containing CBG (at 4x and 8x MIC) set up in 96-well microtiter plates, 36 wells/concentration. Plates were incubated at 37°C for 4 days. Alternatively, bacteria were treated with a 2-fold series of CBG concentrations spanning the MIC. Bacteria growing at the maximum sub-MIC concentration were repeatedly passaged in a similar series of CBG concentrations by 1000-fold dilution every 24 hours. Five CBG dilution series were performed simultaneously and the cells were passaged for 15 days.

### General molecular techniques

DNA manipulations were performed as previously described ^37^. CaCl_2_ chemically-competent ML35 cells were transformed with pBR322 encoding a periplasmic β-lactamase.

### Biofilm formation assays

Biofilm formation was performed in polystyrene 96-well plates in Tryptic Soy Broth (TSB) with 1% glucose and detected by the crystal violet method as previously described^38^. For MBEC determination, biofilms were allowed to form for 24 hours prior to washing off planktonic cultures with sterile de-ionized water then treatment with CBG at varying concentrations.

Quantification of biofilm was performed as noted above.

### Membrane integrity assays

DiSC_3_(5) assay was performed in *S. aureus* as previously described^39^. To determine outer membrane and inner membrane activity of CBG against Gram-negative bacteria, we performed β-lactamase and β-galactosidase assays, respectively. Overnight cultures of ML35 pBR322 in TSB with 50 μg/mL ampicillin were 100-fold diluted in fresh pre-warmed TSB and incubated at 37°C at 220 rpm. Logarithmic phase cells were collected, washed twice in PBS and then resuspended in PBS at a final OD_600_ of 0.01. Nitrocefin (30 μM) or ONPG (1.5 mM) - probes for β-lactamase and β-galactosidase, respectively (final concentration) - were added to the bacterial suspension and immediately aliquoted to dilution series of CBG and/or PMB at 100 μL final volume. Plates were incubated at 37°C and monitored kinetically for color change at 492 and 405 nm (for nitrocefin and ONPG hydrolysis, respectively). Adequate no drug, no probe and/or cell-free controls were included.

### Hemolysis assay

Hemolysis assay using red blood cells (defibrinated sheep blood, Thermo Fisher Scientific) was performed as previously described^40^.

### Statistical analyses

Statistical analyses were conducted with GraphPad Prism 5.0 and is indicated for each assay in the figure caption. All results are shown as mean ±SEM unless otherwise stated. In the case of MIC and checkerboard assays, the experiments were repeated at least three independent times and the experiment showing the most conservative effects (if applicable) was shown.

### Synthesis of Cannabigerol

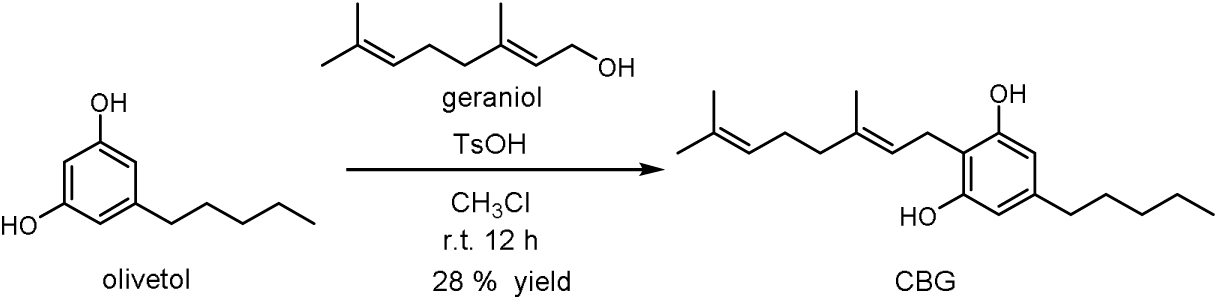

CBG was synthesized using a reported procedure^41^. To a 25 mL round-bottomed flask containing a magnetic stir were added olivetol (108 mg, 0.6 mmol), chloroform (5 mL), geraniol (174 µL, 1.0 mmo), *p*-toluene sulfonic acid monohydrate (19 mg, 0.1 mmol). The flask was covered with aluminum foil and the reaction was stirred at room temperature in the dark for 12 hours at which point TLC analysis indicated complete consumption of the olivetol substrate. To the reaction was added aqueous saturated NaHCO_3_ (5 mL). The organic phase was removed and washed with water (5 mL). The aqueous layer was extracted with additional chloroform (5 mL) and the combined organic extracts were dried over MgSO_4_ and concentrated *en vacuo*. The crude residue was purified via flash column chromatography on silica gel using gradient elution with hexanes and ethyl acetate. CBG was isolated as an off white powder in 28 % yield (54 mg, 0.17 mmol). Spectral data can be found in Supplementary Figs. 12-14.

^**1**^**H NMR** (700 MHz, CDCl_3_) δ 6.25 (s, 2H), 5.28 (tq, J = 7.1, 1.3 Hz, 1H), 5.09 – 5.04 (m, 3H), 3.40 (d, J = 7.1 Hz, 2H), 2.49 – 2.43 (m, 2H), 2.14 – 2.04 (m, 4H), 1.82 (s, 3H), 1.68 (s, 3H), 1.60 (s, 3H), 1.58 – 1.54 (m, 2H), 1.36 – 1.28 (m, 4H), 0.89 (t, J = 7.0 Hz, 3H). ^**13**^**C NMR** (176 MHz, CDCl3) δ 154.92, 142.89, 139.13, 132.19, 123.89, 121.84, 110.73, 108.52, 39.83, 35.65, 31.63, 30.93, 26.52, 25.81, 22.68, 22.40, 17.83, 16.32, 14.16. HRMS (ESI) *m/z*: 315.2329 calcd for C_21_H_31_O_2_ [M-H]; 315.24490 obsd.

### Mouse infection models

Animal experiments were conducted according to guidelines set by the Canadian Council on Animal Care using protocols approved by the Animal Review Ethics Board at McMaster University under Animal Use Protocol #17-03-10. Before infection, mice were relocated at random from a housing cage to treatment or control cages. No animals were excluded from analyses, and blinding was considered unnecessary. Seven-to nine-week old female CD-1 mice (Envigo) were infected intraperitoneally with 7.5 × 10^7^ CFU of log-phase MRSA strain USA 300 JE2 with 5% porcine mucin. Treatment of 100 mg/kg CBG or a vehicle solution (60% PEG300 and 5% DMSO) were administered intraperitoneally immediately post-infection. Mice were euthanized and tissues collected into phosphate buffered saline (PBS) at necropsy. Organs were homogenized using a high-throughput tissue homogenizer, serially diluted in PBS, and plated onto solid LB. Plates were incubated overnight at 37°C and colonies were quantified to determine organ load.

For Table of Contents Use Only

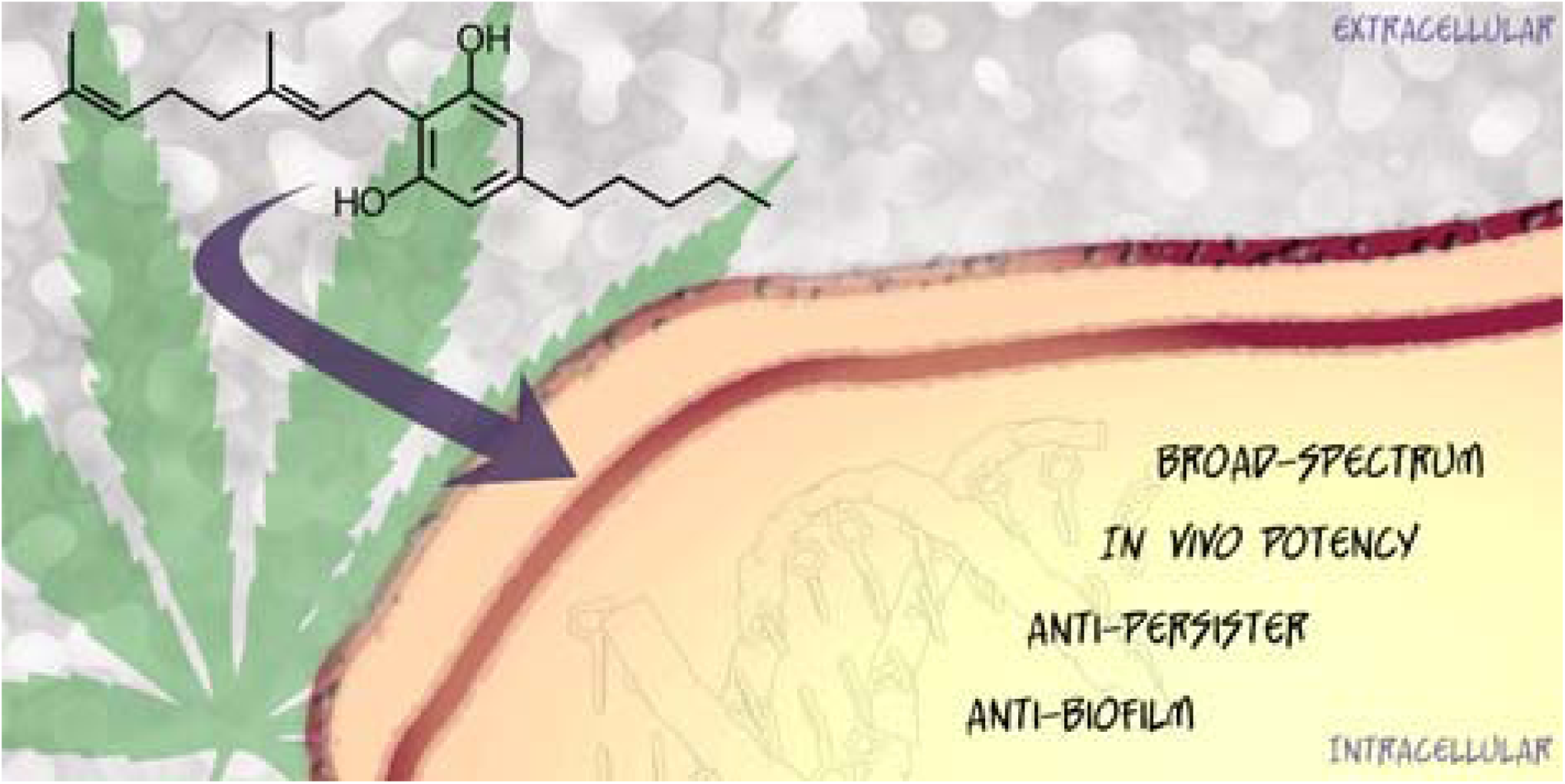

