## Supplementary information for "Uncovering the hidden antibiotic potential of Cannabis"

† These authors contributed equally

**Supplementary Table 1.** Antimicrobial activity of cannabinoid analogs against MRSA USA300

| Entry | Compound Name (Abbreviation) | | Structure | MIC  (µg/mL) |
| --- | --- | --- | --- | --- |
| 1 | cannabigerol | (CBG) | 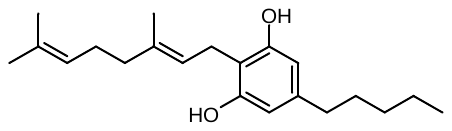 | 2 |
| 2 | cannabidiol | (CBD) | 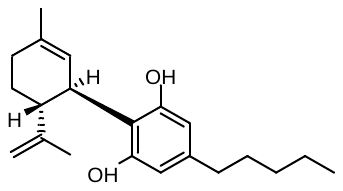 | 2 |
| 3 | cannabinol | (CBN) | 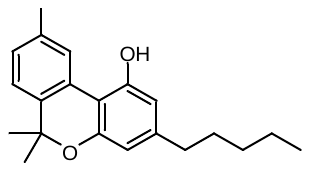 | 2 |
| 4 | cannabichromene | (CBC) | 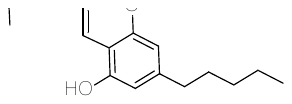 | 8 |
| 5 | cannabichromenic acid | (CBCA) | 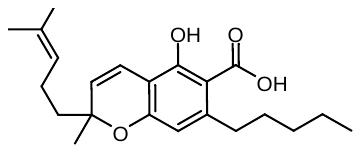 | 2 |
| 6 | (-)Δ^9^-tetrahydrocannabinol | (THC) | 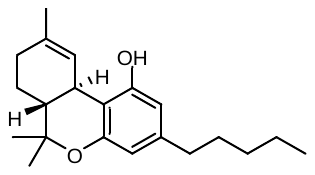 | 2 |
| 7 | (-)Δ^8^-tetrahydrocannabinol | (-Δ8THC) | 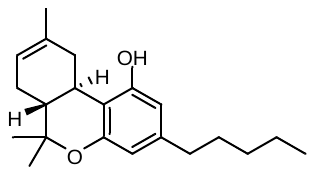 | 2 |
| 8 | exo-tetrahydrocannabinol | (exo-THC) | 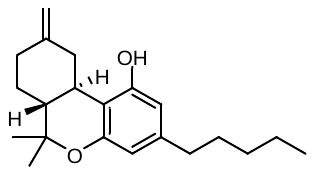 | 2 |
| 9 | Δ^9^-tetrahydrocannabinolic  acid A | (THCAA) | 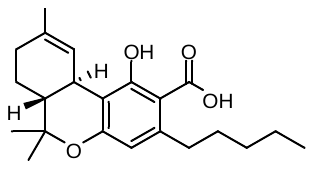 | 4 |
| 10 | Δ^9^-tetrahydrocannabivarin | (THCV) | 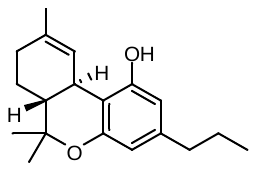 | 4 |
| 11 | cannabigerolic acid | (CBGA) | 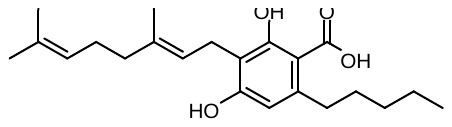 | 4 |
| 12 | cannabidivarin | (CBDV) | 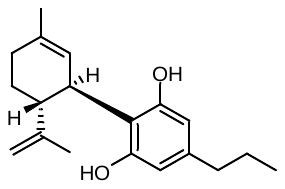 | 8 |
| 13 | cannabidiolic acid | (CBDA) | 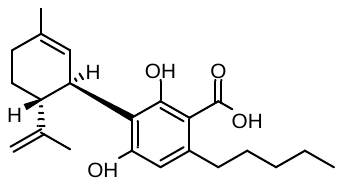 | 16 |
| 14 | tetrahydrocannabivarinic acid | (THCVA) | 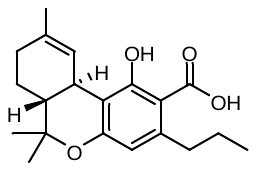 | 16 |
| 15 | cannabidivarinic acid | (CBDVA) | 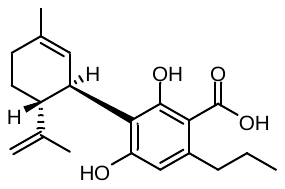 | 32 |
| 16 | (±) 11-nor-9-carboxy-Δ^9^-THC |  | 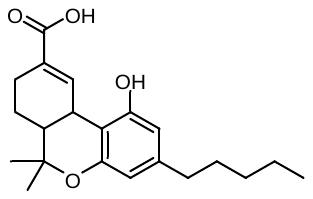 | >32 |
| 17 | (±) 11-hydroxy-Δ^9^-THC |  | 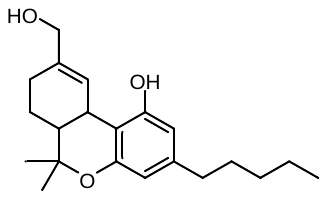 | >32 |
| 18 | cannabicyclol | (CBL) | 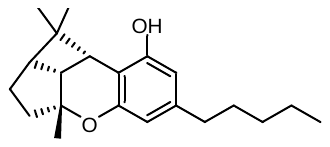 | >32 |

**Supplementary Table 2.** MRSA transposon mutants sensitized to sub-lethal concentrations of CBG.

| Gene | Function | Transposon |
| --- | --- | --- |
| aroC | chorismate synthase | SAUSA300_1357 |
|  | putative endoribonuclease L-PSP | SAUSA300_0474 |
| estA | tributyrin esterase | SAUSA300_2564 |
| - | hypothetical protein | SAUSA300_0553 |
| - | BioY family protein | SAUSA300_2233 |
| atpA | ATP synthase F1, alpha subunit | SAUSA300_2060 |
|  | conserved hypothetical protein | SAUSA300_1780 |
| graS | sensor histidine kinase | SAUSA300_0646 |
| tcaB | teicoplanin resistance associated membrane protein TcaB protein | SAUSA300_2301 |
| - | hypothetical protein | SAUSA300_2330 |
|  | conserved hypothetical protein | SAUSA300_0199 |
|  | accessory secretory protein Asp1 | SAUSA300_2587 |
|  | putative hemolysin III | SAUSA300_2129 |
| - | hypothetical protein (pyrazinamidase/nicotinamidase pncA) | SAUSA300_1899 |
|  | conserved hypothetical protein | SAUSA300_2212 |
| thiE | thiamine-phosphate pyrophosphorylase | SAUSA300_2047 |
|  | conserved hypothetical protein | SAUSA300_0847 |
| cap5B | capsular polysaccharide biosynthesis protein Cap5B | SAUSA300_0153 |
| qoxC | quinol oxidase, subunit III | SAUSA300_0961 |
| ribBA | riboflavin biosynthesis protein | SAUSA300_1713 |
| sdhA | succinate dehydrogenase, flavoprotein subunit | SAUSA300_1047 |
| hemB | delta-aminolevulinic acid dehydratase | SAUSA300_1615 |
|  | conserved hypothetical protein | SAUSA300_1294 |
| - | TENA/THI-4 family protein | SAUSA300_2050 |
| topB | DNA topoisomerase III | SAUSA300_2208 |
| - | pyruvate ferredoxin oxidoreductase, alpha subunit | SAUSA300_1182 |
| - | pyridoxal biosynthesis lyase PdxS | SAUSA300_0504 |
| sdhB | succinate dehydrogenase iron-sulfur subunit | SAUSA300_1048 |
| sgtB | monofunctional glycosyltransferase | SAUSA300_1855 |
|  | glycosyl transferase, group 1 family protein | SAUSA300_0550 |
|  | putative membrane protein | SAUSA300_0917 |
| bioD | dethiobiotin synthase | SAUSA300_2373 |
| mqo | malate:quinone oxidoreductase | SAUSA300_2312 |
|  | cation efflux family protein | SAUSA300_2099 |
|  | putative lipase/esterase | SAUSA300_0641 |
| - | drug transporter | SAUSA300_2451 |
| sucD | succinyl-CoA synthetase subunit alpha | SAUSA300_1139 |
| lspA | lipoprotein signal peptidase | SAUSA300_1089 |
| pckA | phosphoenolpyruvate carboxykinase | SAUSA300_1731 |
| msrA | methionine sulfoxide reductase A | SAUSA300_1256 |
| - | PTS system, galactitol-specific enzyme II, B component | SAUSA300_0240 |

**Supplementary Table 3.** Bacterial strains and plasmids used in this study.

| Bacterial Strain | Description | Reference |
| --- | --- | --- |
| *S. aureus*  USA300  NTML  *B. subtilis* 168  CRISPRi collection  *P. aeruginosa*  PAO1  *E. coli*  ML35    ML35pBR322  K-12 BW25113  *A. baumannii*  ATCC19606  ATCC19606-LOS^-^ | USA300 LAC isolated in 2002 from a skin and soft tissue infection of an inmate in the Los Angeles County Jail in California, USA.; hypervirulent community-associated MRSA; cured of antibiotic resistance plasmid; also known as JE2; parent of the NTML.    Nebraska Transposon Mutant Library Screening Array; 1920 *S. aureus* subsp. aureus USA300 JE2, transposon (Tn) mutants arrayed in five 384-well microtiter plates. Ery^R^  *B. subtilis* CRISPRi essential gene knockdown strain collection  Clinical isolate  *lacZ^+^Y^-^I^-^, E. coli* with constitutive expression of β-galactosidase but lacking the lactose permease  ML35 with periplasmic β-lactamase  *A. baumannii* lacking lipooligosaccharides (LOS) | Laboratory stock*  NARSA*  BGSC**  ^1^  ^2^  ^3^  This study  Lab stock  ATCC  ^4^ |
| Bacterial Plasmid | Description | Reference |
| pBR322 | Promoterless bioluminescent reporter plasmid encoding *luxABCDE*, Amp^R^ Cm^R^ | ^5^ |

* Provided by the Network on Antimicrobial Resistance in *Staphylococcus aureus* (NARSA) for distribution by BEI Resources, NIAID, NIH.

** Bacillus Genetic Stock Center

Amp, ampicillin; Ery, erythromycin; Cm, chloramphenicol; Spec, spectinomycin; Strep, streptomycin

Strains and plasmids constructed in this study are available from the authors upon request.

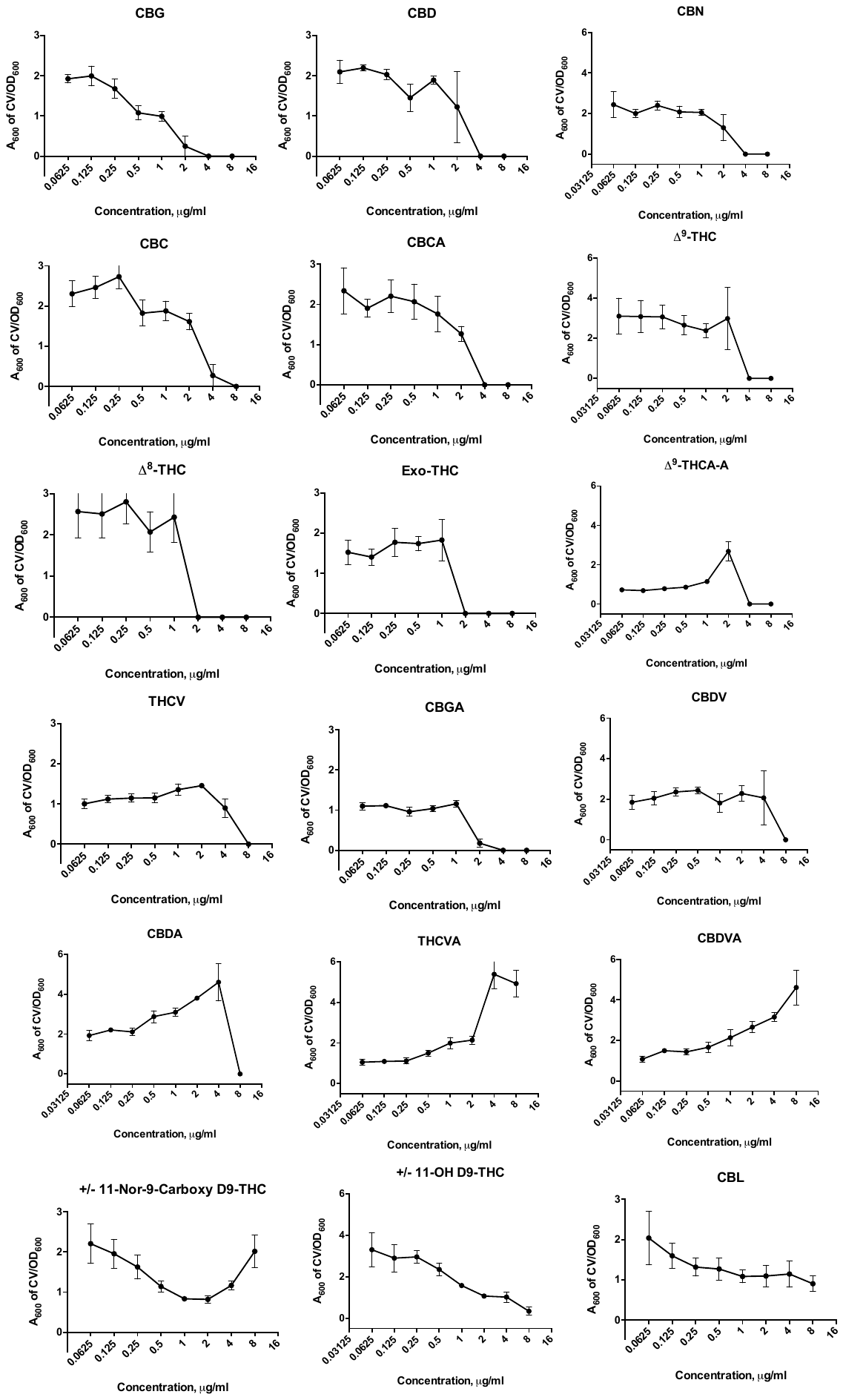

**Supplementary Fig. 1.** Effect of cannabinoids on biofilm formation of MRSA USA300. Shown is the effect of increasing concentrations of the cannabinoids on MRSA biofilm formation. The average A_600nm_ measurements of crystal violet stained biofilms normalized by the OD_600_ of planktonic cells are shown with error bars representing S.E.M. (n = 4). Some molecules seem to lead to increases in A_600_ of CV/OD_600_ at certain concentrations, which is consistent with the observation that many antibacterial agents stimulate biofilm formation at subinhibitory concentrations^6-7^.

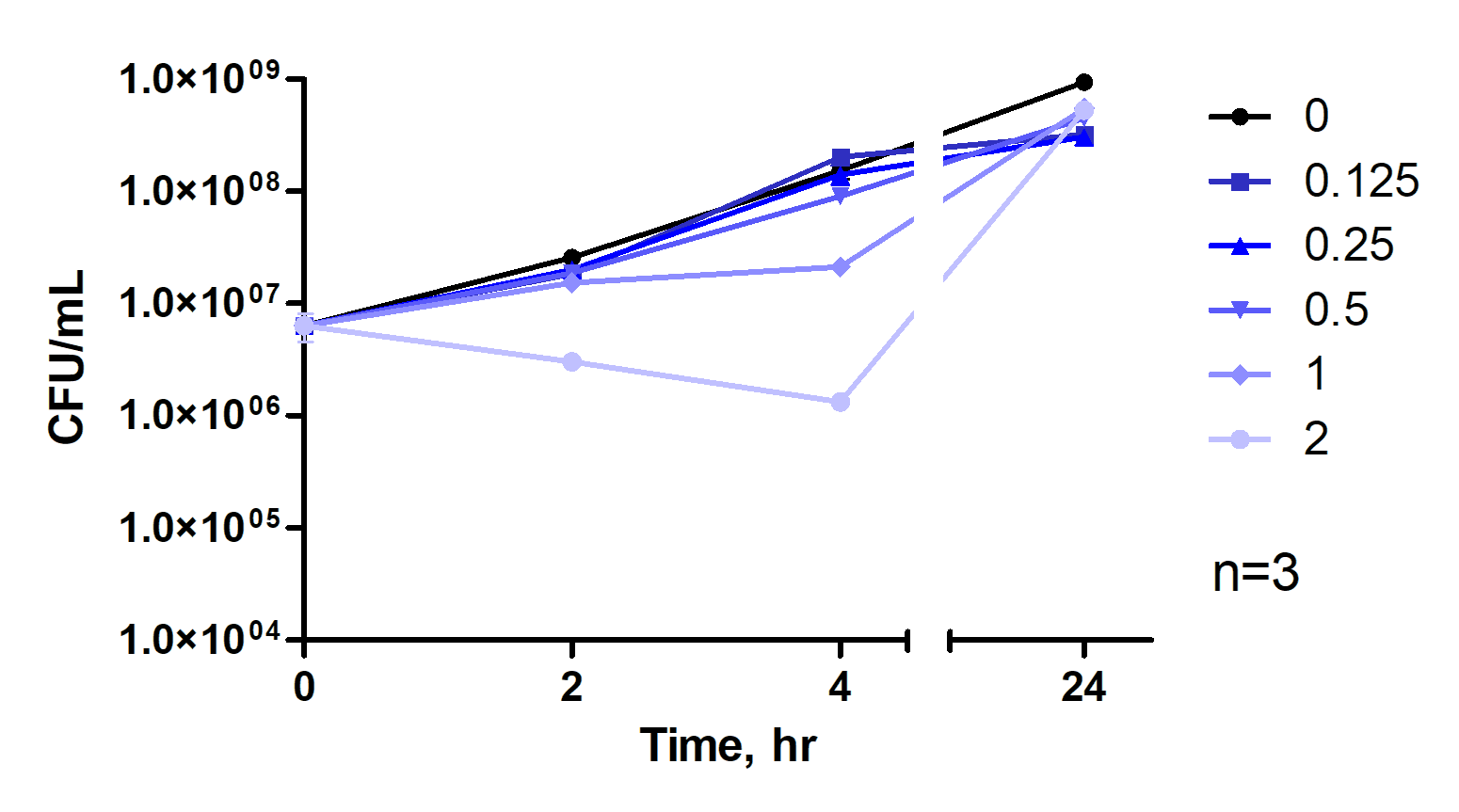

Supplementary Fig. 2. Kill curve performed under conditions mimicking those of the biofilm formation assays. For these experiments, CBG was added in final concentrations of 0, 0.125, 0.25, 0.5, 1 and 2 µg/mL to bacterial cultures (~5x10^6^ CFU/ml), and aliquots were taken after 0, 2, 4, and 24 h. These were serially diluted with sterile PBS and plated on agar plates. After incubation for 24 h at 37 °C, colonies were counted.

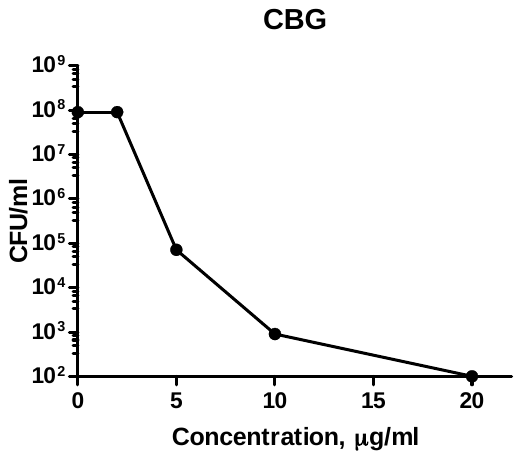

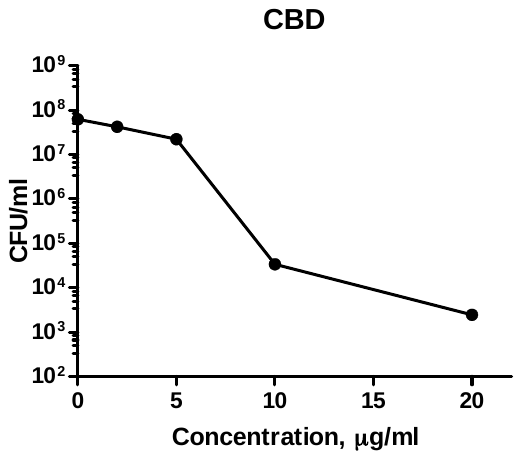

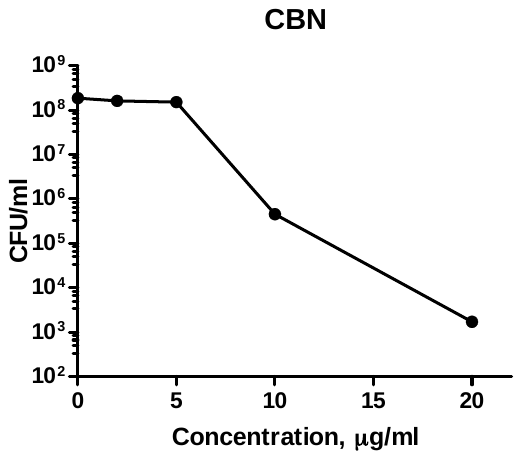

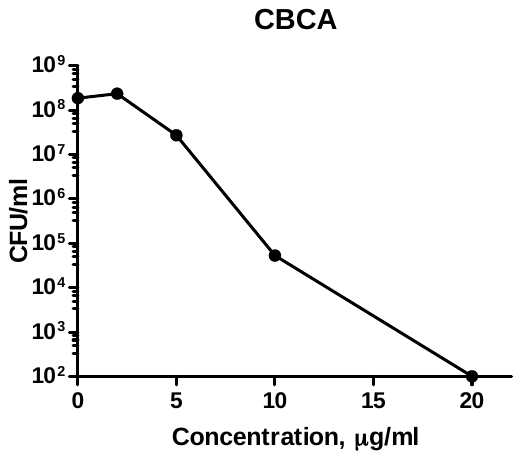

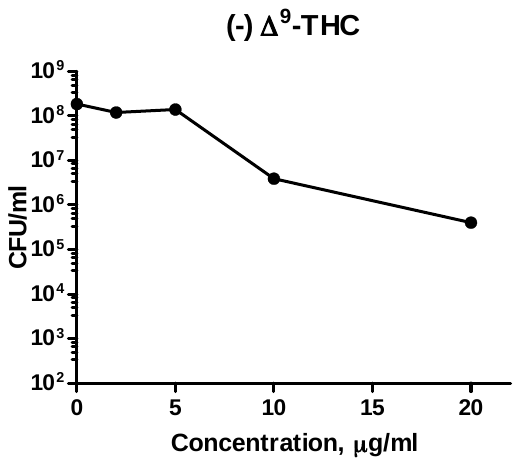

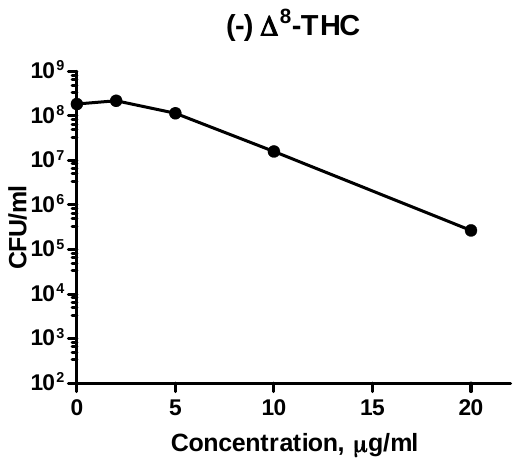

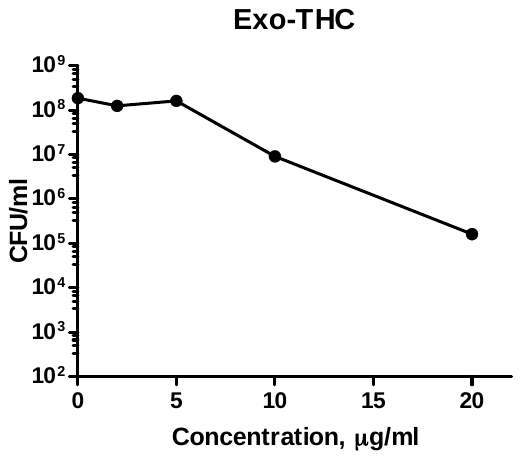

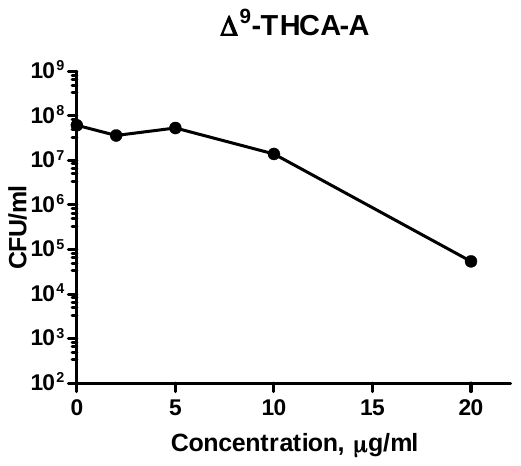

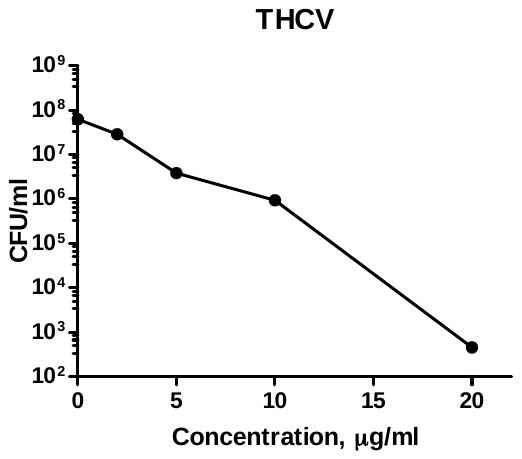

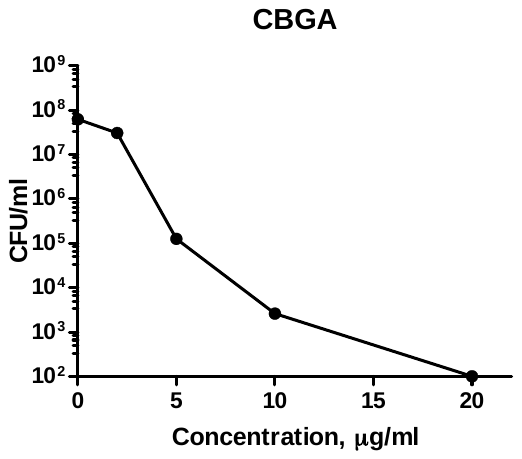

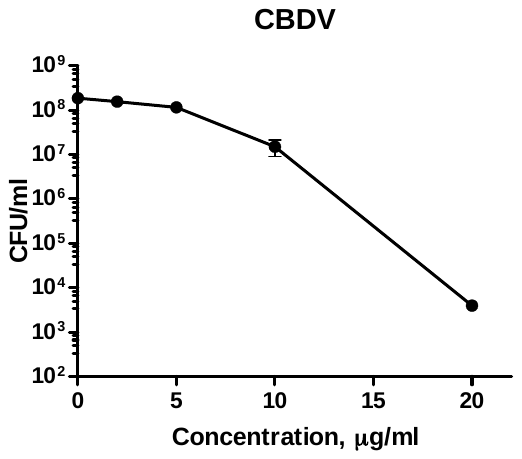

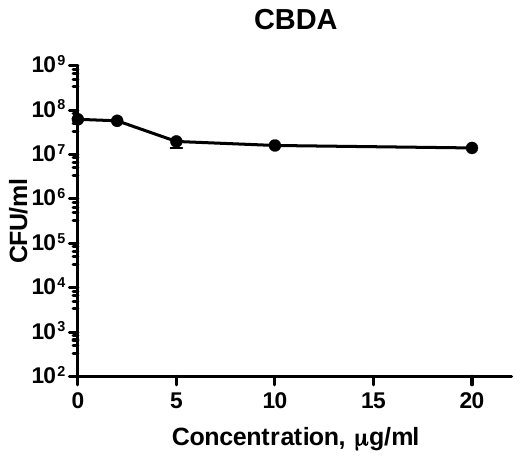

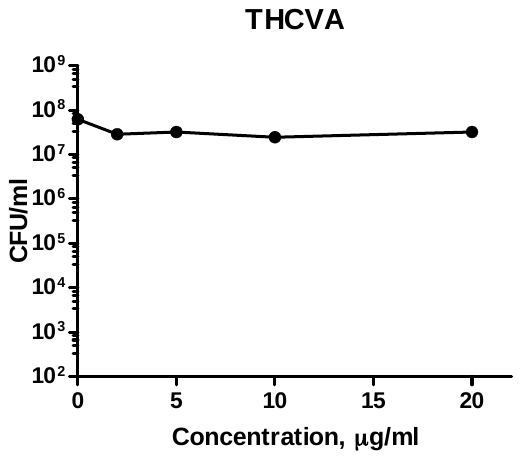

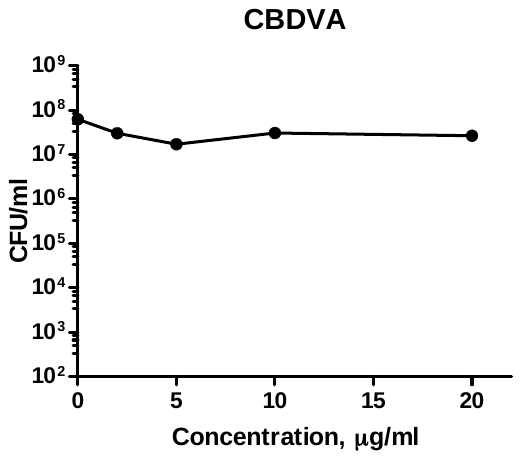

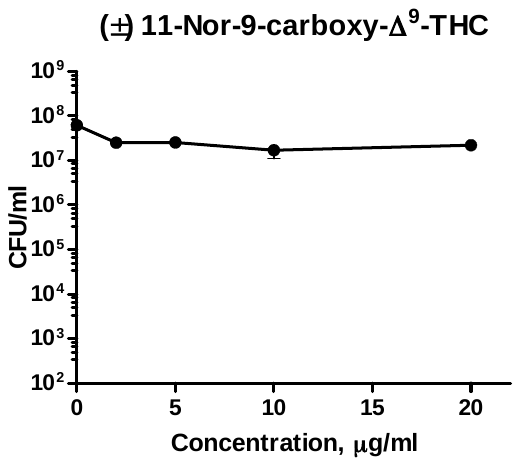

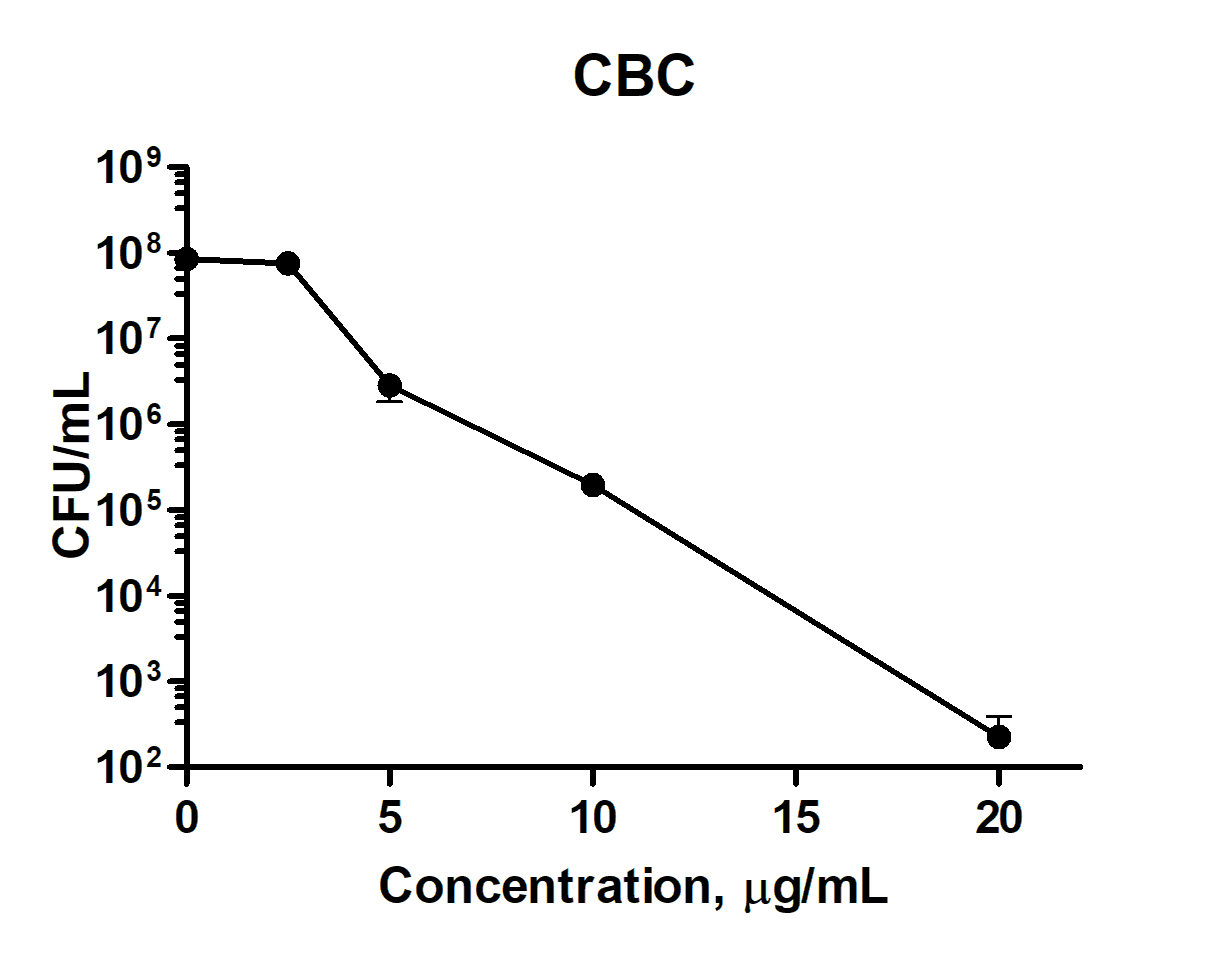

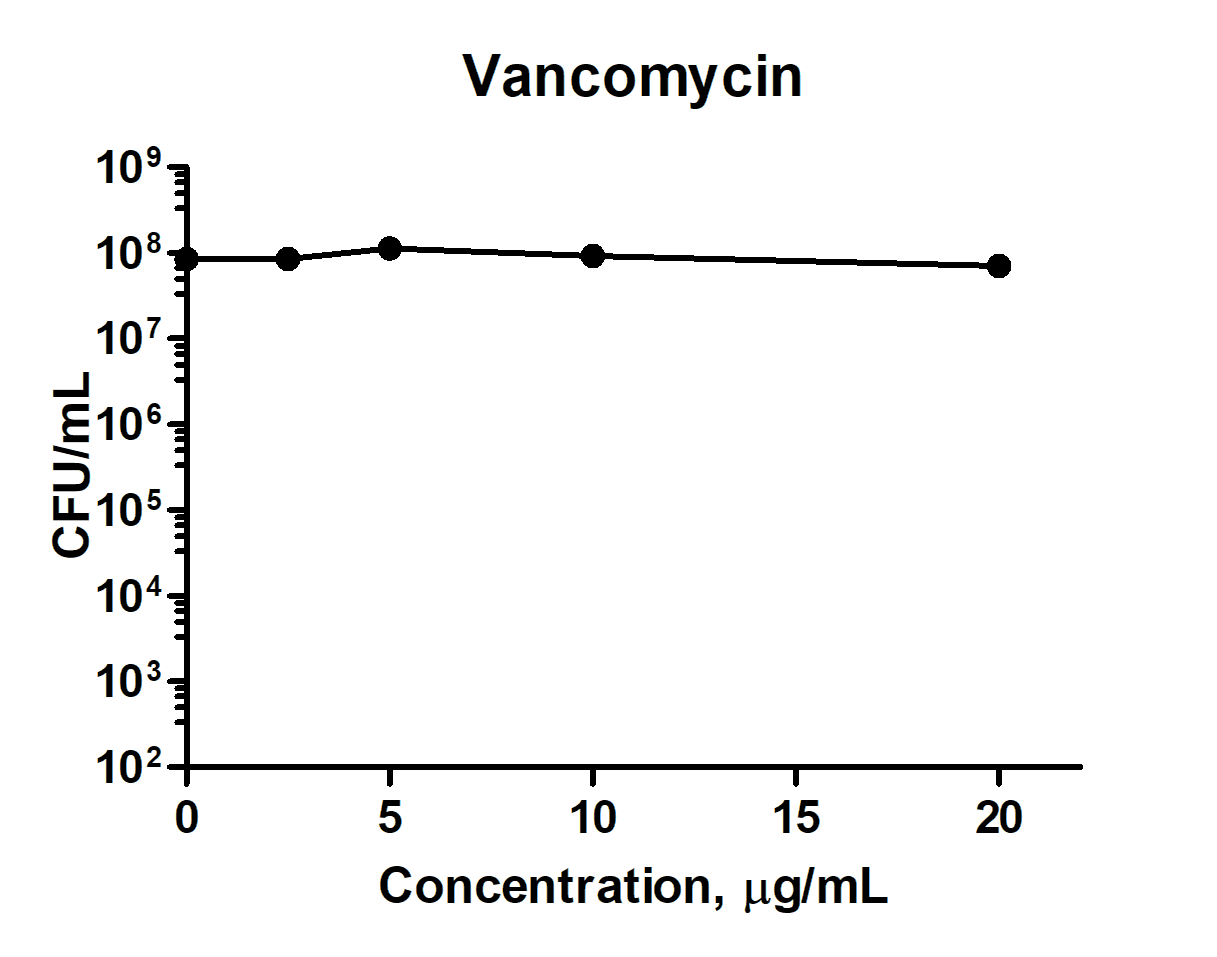

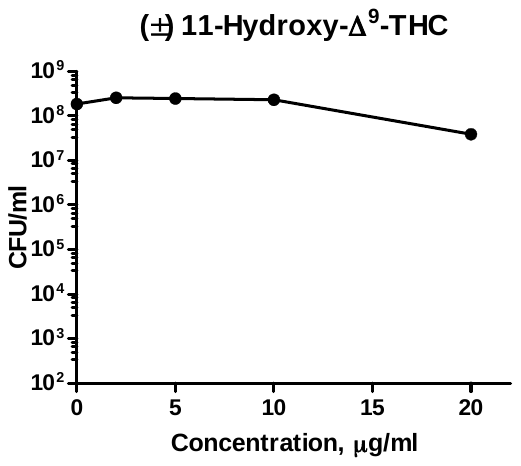

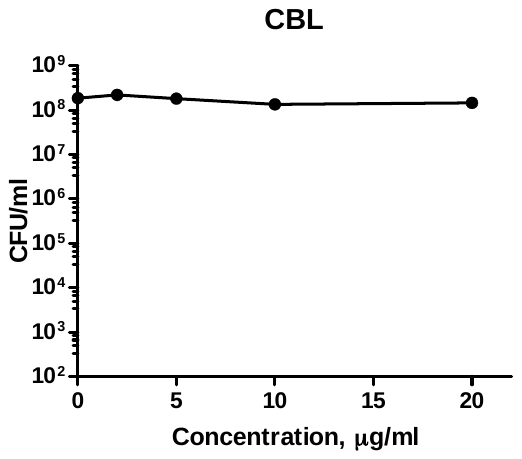

**Supplementary Fig. 3**. Killing of *S. aureus* USA300 persisters by various cannabinoids detected after 1 hour of treatment shown as mean ±S.E.M.

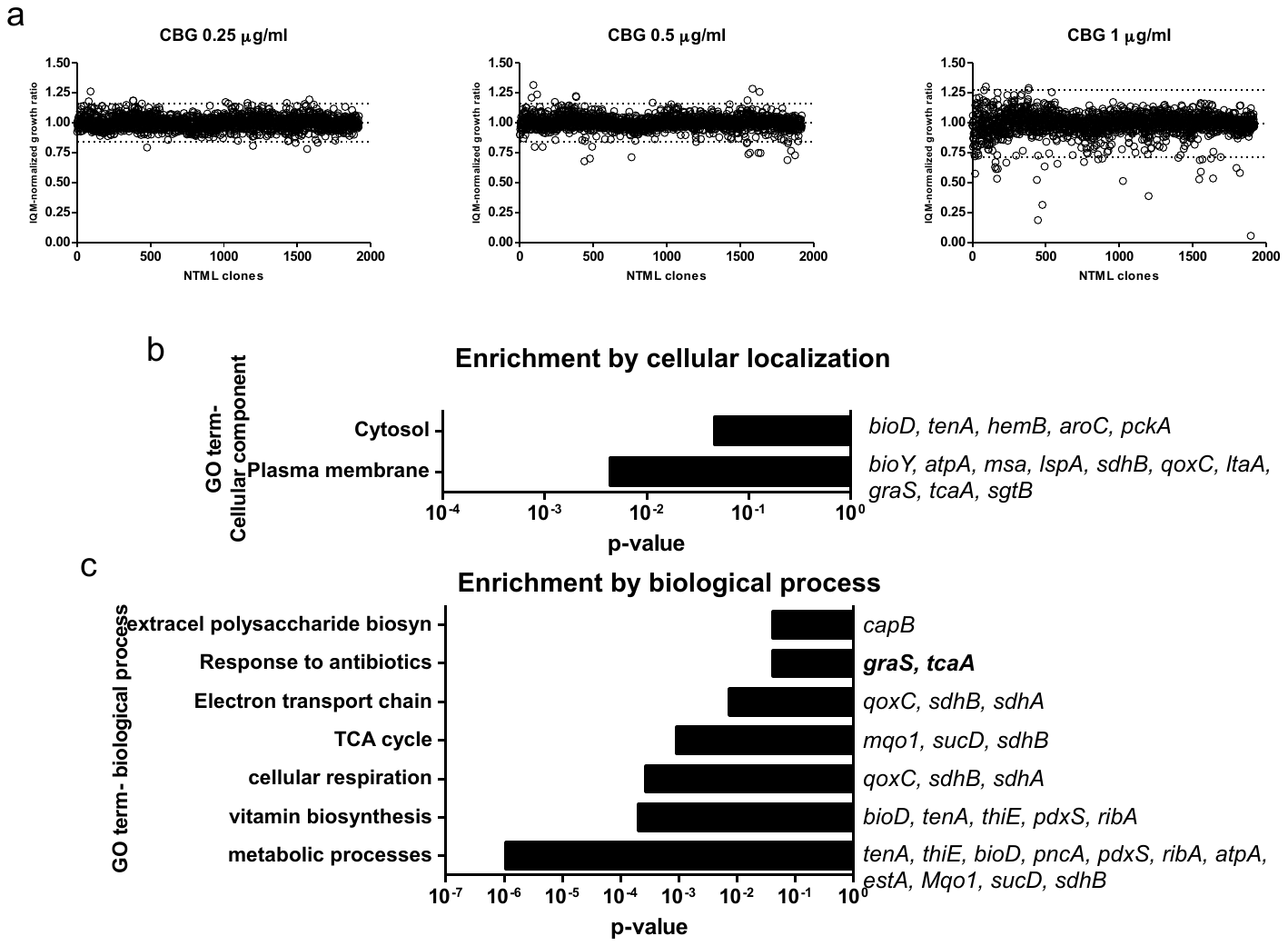

**Supplementary Fig. 4**. Chemical genomic studies using the Nebraska Transposon Mutant Library. **a**, Chemical genomic screen of the transposon library at various sub-lethal concentrations of CBG. **b**, Enrichment by cellular localization, whereby transposon mutants were classified based on gene ontology (GO). Enrichment was based on functional overrepresentation of the mutants resulting in sensitivity to CBG, using a Fisher’s exact test to calculate p-value. **c**, Enrichment by biological processes, whereby transposon mutants were classified based on gene ontology (GO). Enrichment was based on functional overrepresentation of the mutants resulting in sensitivity to CBG, using a Fisher’s exact test to calculate p-value.

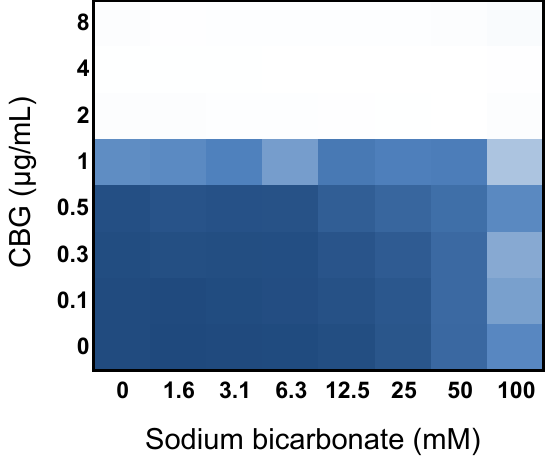

**Supplementary Fig. 5.** Combination of CBG with sodium bicarbonate against MRSA USA300. The extent of inhibition is shown as a heat plot, such that the darkest blue color represents full bacterial growth.

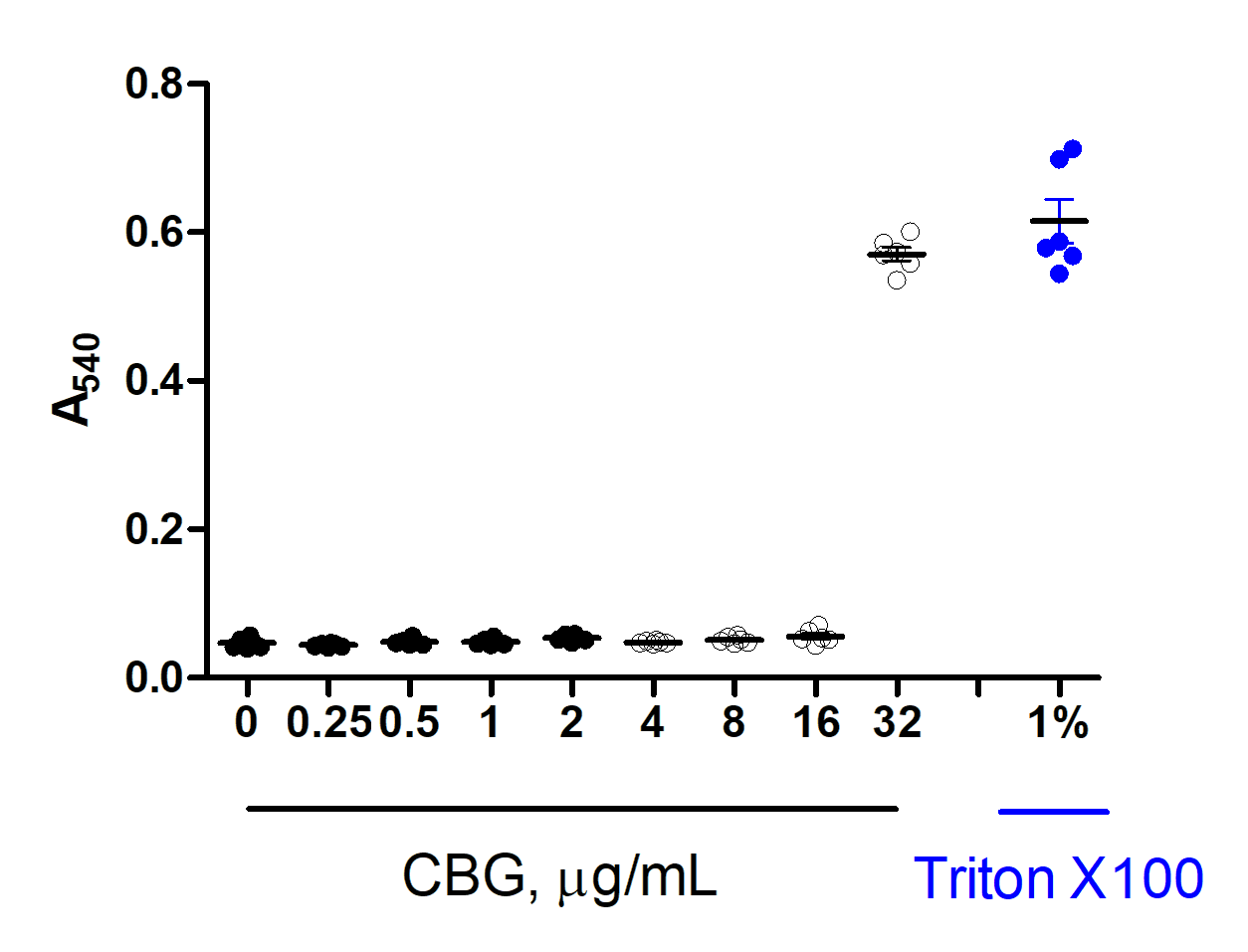

**Supplementary Fig. 6.** Hemolytic activity of CBG over various concentrations. 1% Triton X- 100 was used as a positive control.

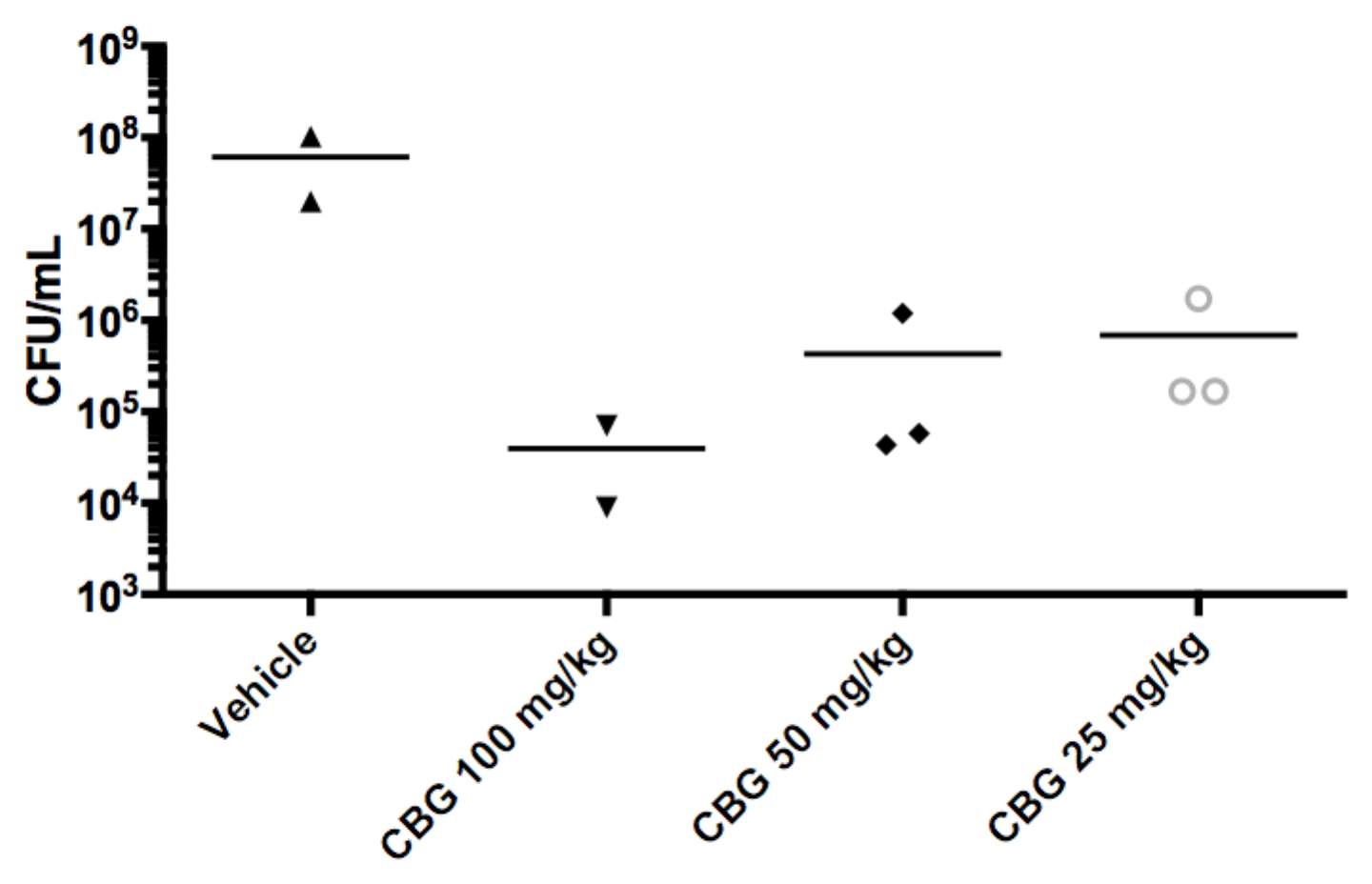

Supplementary Fig. 7. *In vivo* dose-dependence of CBG in treating a systemic mouse model of *S. aureus* infection. CD-1 mice were infected intraperitoneally with 7.5 x 10^7^ CFU of log-phase MRSA strain USA 300 JE2 with 5% porcine mucin. Treatment of varying doses of CBG or a vehicle solution (60% PEG300 and 5% DMSO) were administered intraperitoneally immediately post-infection. Mice were euthanized 8 hours post-infection and tissues collected. Spleens were homogenized, plated and colonies quantified to determine organ load.

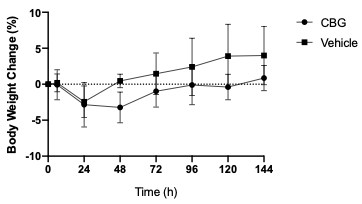

Supplementary Fig. 8. Percent changes in body weight of mice at various time points following treatment with 100 mg/kg CBG (circle, n=5, i.p.) or a vehicle control (square, n=4, i.p.).

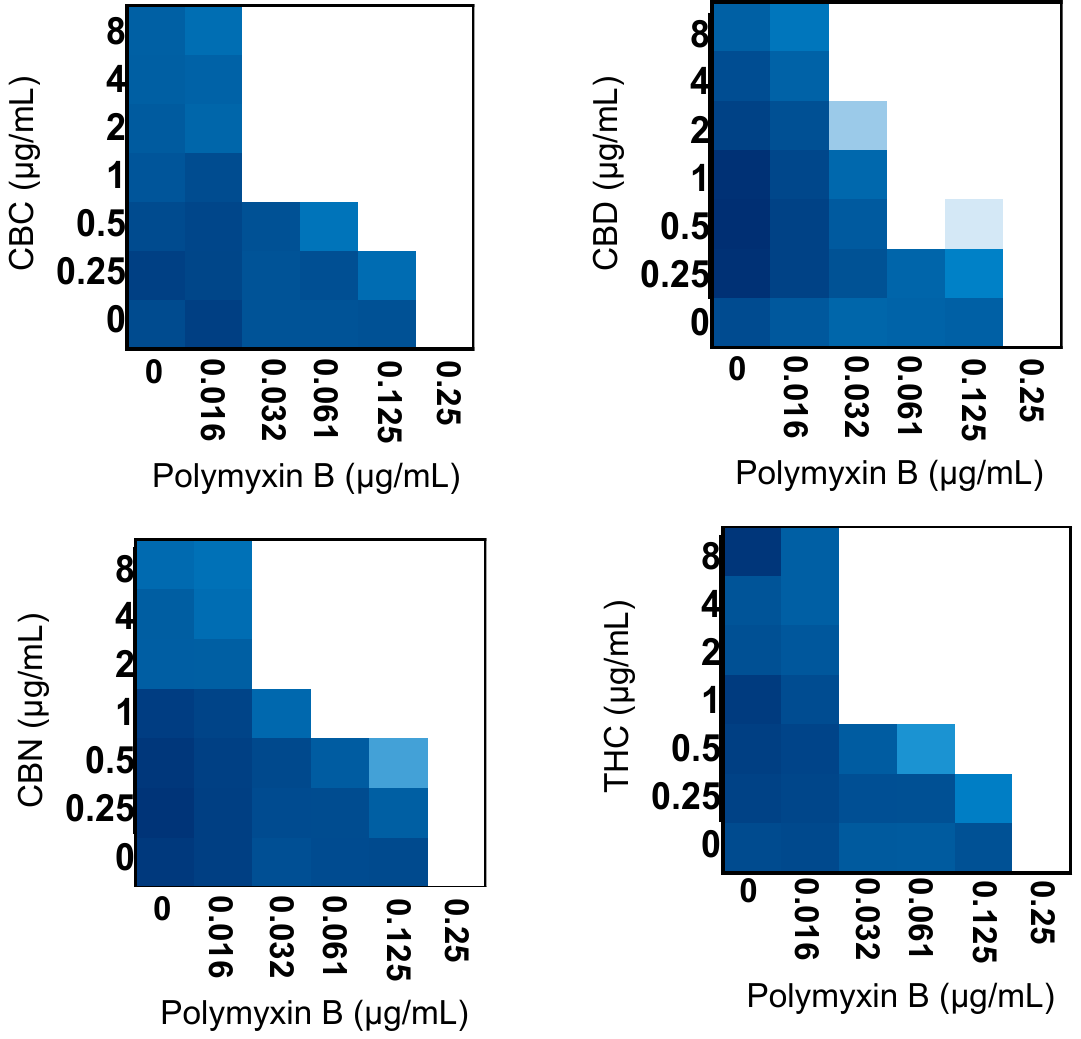

**Supplementary Fig. 9.** Checkerboard analysis of CBC, CBD, CBN and THC in combination with polymyxin B against *E. coli* (K-12 BW25113). The extent of inhibition is shown as a heat plot, such that the darkest blue color represents full bacterial growth.

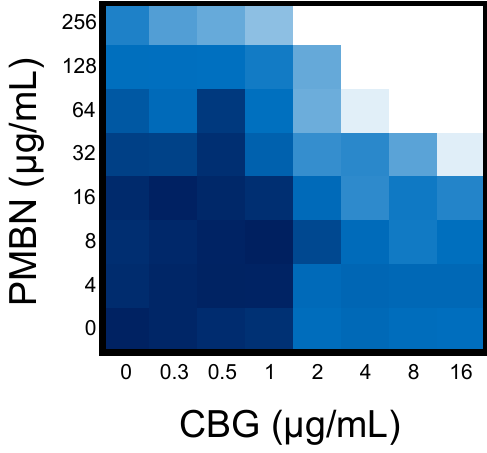

Supplementary Fig. 10. Checkerboard analysis of CBG and polymyxin B nonapeptide (PMBN) against E. coli (K-12 BW25113). The extent of inhibition is shown as a heat plot, such that the darkest blue color represents full bacterial growth.

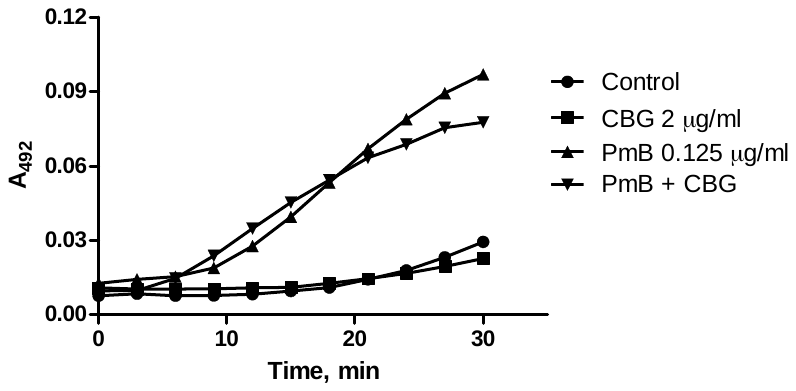

**Supplementary Fig. 11.** CBG is not active against the OM of *E. coli*, as measured by hydrolysis (absorbance at 492 nm) of nitrocefin upon permeation across the OM of an *E. coli* expressing a periplasmic β-lactamase.

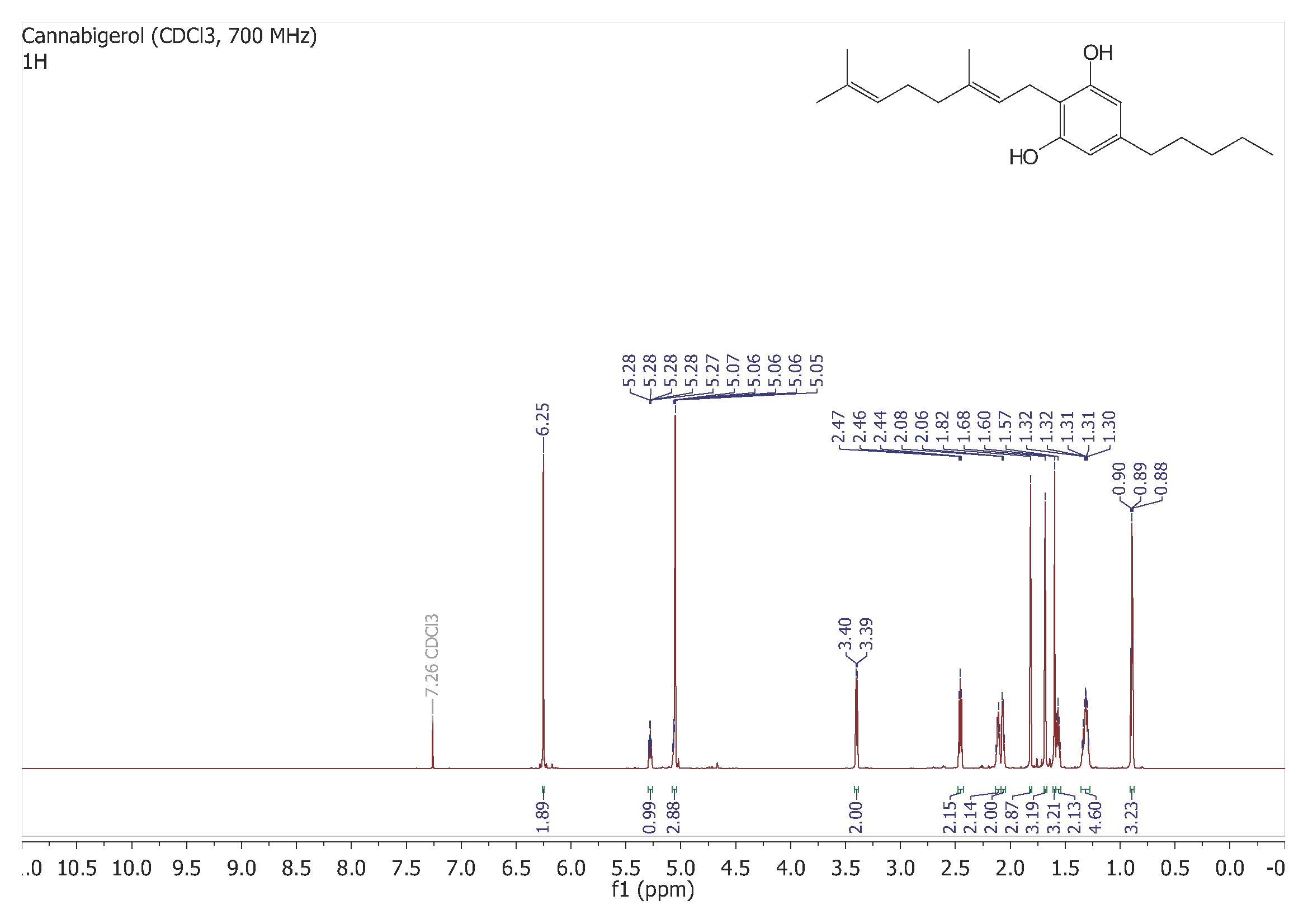

**Supplementary Fig. 12.** ^1^H NMR spectra of CBG. Chemical shifts in the ^1^H NMR spectrum is reported in parts per million (ppm) relative to tetramethylsilane (TMS), with calibration of the residual solvent peaks according to values reported by Gottlieb *et al.* (chloroform: δ_H_ 7.26)^8^. The ^1^H NMR spectrum was acquired at 700 MHz with a default digital resolution (Brüker parameter: FIDRES) of 0.15 Hz/point and coupling constants reported herein therefore have uncertainties of ±0.30 Hz.

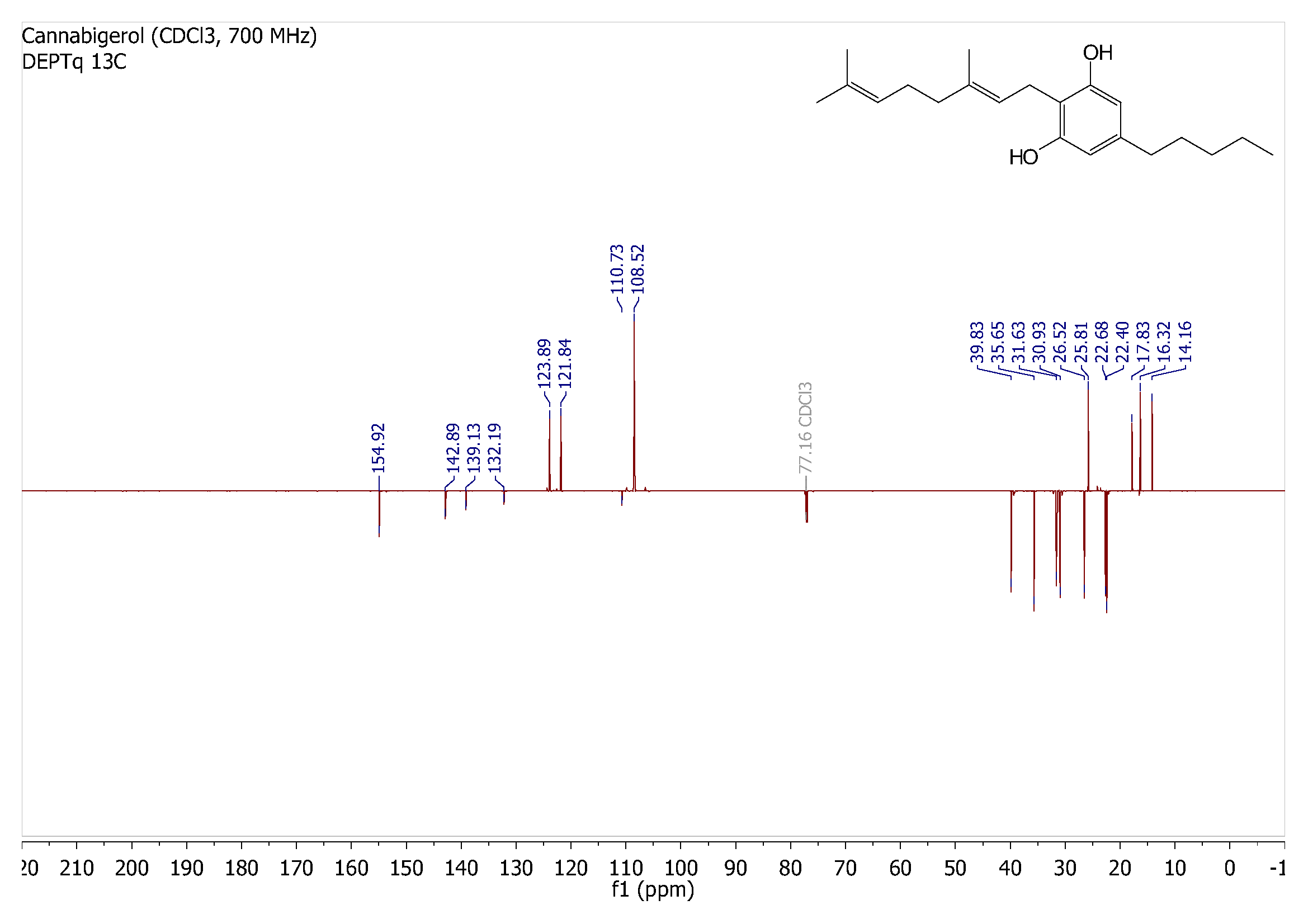

**Supplementary Fig. 13.** ^13^C NMR spectra of CBG. Chemical shifts in the ^13^C NMR spectrum are reported in parts per million (ppm) relative to tetramethylsilane (TMS), with calibration of the residual solvent peaks according to values reported by Gottlieb *et al.* (chloroform: δ_C_ 77.16)^8^. The ^13^C NMR spectrum was acquired at 176 MHz.

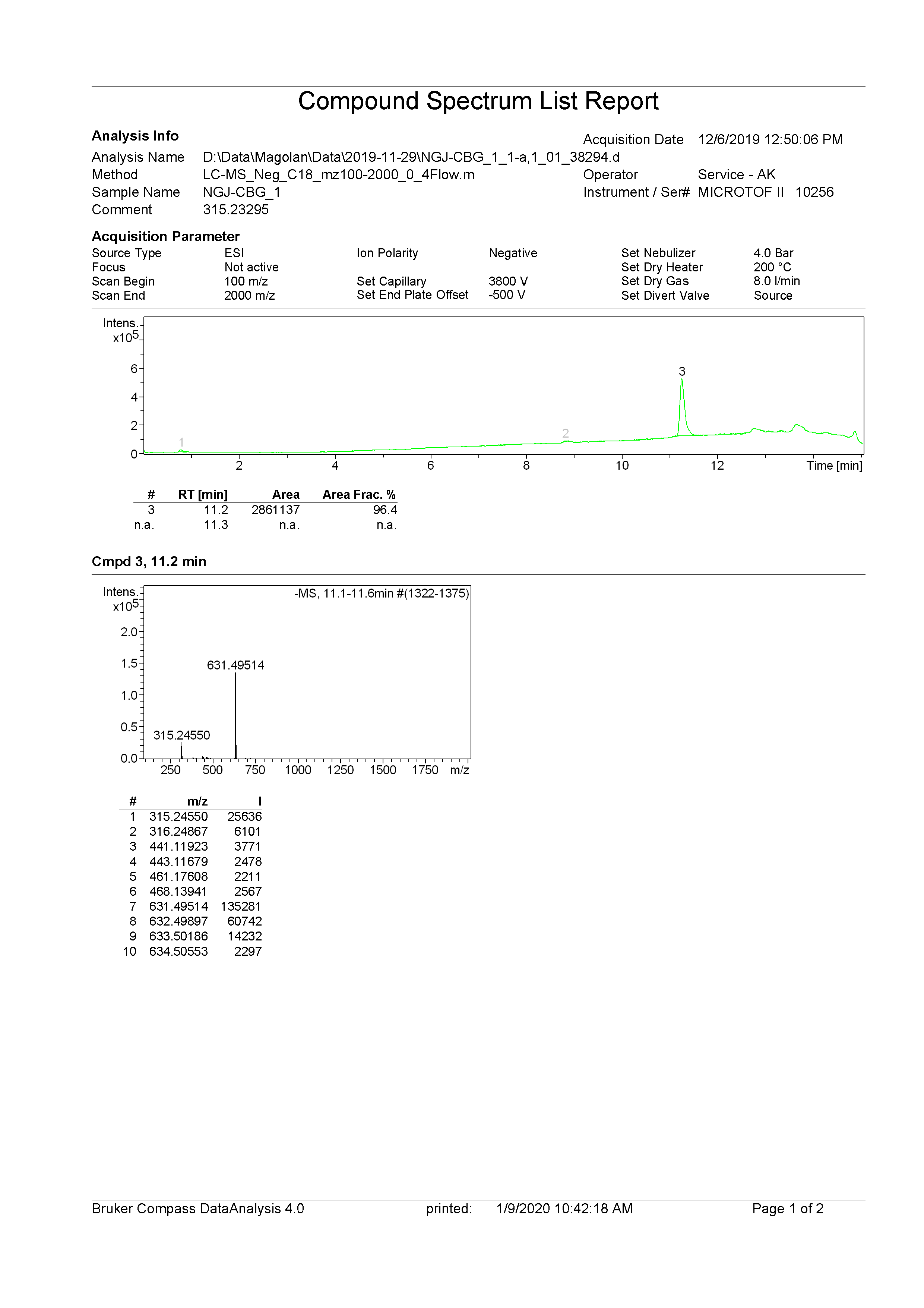

**Supplementary Fig. 14.** High Resolution Mass Spectrum of CBG. The LC-MS purity analysis determined CBG to be 96.4% pure.
